# Evaluation of Drug/Ligand Binding Constants for Human Serum Albumin Using Differential Scanning Calorimetry

**DOI:** 10.1101/2021.01.22.427858

**Authors:** Matthew W. Eskew, Albert S. Benight

## Abstract

This paper reports utilization of differential scanning calorimetry measurements to evaluate binding constants for Human Serum Albumin of 28 different drug ligands. Protein/ligand mixtures were prepared at various ligand concentrations and subjected to thermal denaturation analysis by calorimetry. From the measurements, the melting temperature, *T_m_*, and free-energy *ΔG_cal_*(37°*C*) for melting ligand-bound Albumin were evaluated as a function of ligand concentration. Concentration dependent behaviors of *ΔG_cal_*(37°*C*) and *T_m_* derived from protein/ligand mixtures were used to construct dose-response curves. Model fits of dose-response curves yielded quantitative evaluation of the ligand binding constant, *K_D_*, and semi-quantitative estimates of the binding stoichiometry, n. Many of the ligands had known binding affinity for Albumin with binding constants reported in the literature. Evaluated Albumin binding parameters for the ligands impressively agreed with reported literature values determined using other standard experimental methods. These results demonstrated utility of our calorimetry-based process for applications in pre-clinical drug screening.

## INTRODUCTION

Methods for analyzing binding of ligands to proteins are crucial for a number of applications. One of central importance is analysis of drug/ligand binding to Human Serum Albumin (HSA). Assessment of plasma protein binding (PPB) of new drug candidates is an integral component of pre-clinical drug discovery and protein engineering. For the purpose of this study, the terms ligand and drug are used interchangeably to mean binding entities. In essence, all drugs are ligands but not all ligands are necessarily drugs.

At 60% by mass of all plasma proteins HSA is the most prominent.^1–4^ Intrinsic to its biological activities, HSA is directly involved in transport and distribution of therapeutic drugs in the blood. This activity can directly impact desired pharmacological effects of a therapeutic drug. A multiplicity of reactive sites on HSA have been identified, and several with known binding affinities for a variety of different drugs.^5–7^ The pharmacokinetics (ADME properties) of administered therapeutic agents can be greatly impacted by the enormous binding capacity of HSA, and binding of other plasma proteins. Both experimental and computer studies have extensively analyzed site-specific drug binding in HSA.^8–10^ The prominence of HSA among plasma proteins and the central role HSA plays in bioavailability of drugs, makes it an important screening target. Screening of new drug candidates for HSA binding and analysis is central to their pre-clinical evaluation and development.

Supply of new drug candidates exceeds current screening capacity. This is primarily due to the use of ineffective high-throughput screening methods. The result is bottlenecks in the development pipeline and ultimately lower availability of potentially effective drug candidates. Improved methods for rapid, quantitative screening of new drug candidates for their PPB activity, could aid considerably in loosening disruptive bottlenecks and expediting the drug discovery process.^11^ At least 90% of all new drug candidates fail. Improved screening methodologies capable of rapidly evaluating protein binding properties are needed to provide deeper insight into relationships between molecular and chemical characteristics of targets and drug candidates. This information provides the means to establish correlations between specific physical and chemical attributes of drugs and their targets. Such correlations can then be employed in the functional design and optimization of biochemical properties of new drug candidates. These improvements could aid to reduce the failure rate of new drugs and could have significant impact on the early pre-clinical drug discovery process.

Generally, when a drug/ligand recognizes and binds to a protein, depending on the nature of binding (electrostatic, polar, hydrophobic, etc.) it can either stabilize or destabilize the protein with respect to thermal or chemical denaturation. Thus, relative to the unbound protein, when bound the melting temperature (or concentration of denaturant) required to unfold the protein is either increased or decreased. Interactions between circulating drug/ligands in plasma with HSA can affect protein thermodynamic stabilities, thereby influencing the shape of the melting curve or thermogram of drug/ligand-bound HSA.^12–14^ In this sense, thermograms measured by differential scanning calorimetry (DSC) are highly sensitive to drug/ligand binding interactions.

This enhanced sensitivity of ligand-bound HSA compared to free HSA is due to a direct linkage between ligand binding and thermodynamic stability of the protein.^15^ The nature of this linkage is two-fold. Ligand binding to a protein as a function of increased ligand concentration results in increases in melting transition temperatures, *T_M_*, of ligand-bound protein complexes. Not unexpected this observation is simply a well-known manifestation of Le Chatelier’s principle as applied to protein-ligand binding.^16, 17^ That is if a ligand binds preferentially to the native rather than denatured state the thermal stability will be increased. Provided the measured calorimetric enthalpy of the heat-induced denaturation transition of the ligand-bound protein remains constant, as ligand concentration is increased, the observed shift in stability with ligand binding forms the foundation of any standard temperature shift assay.^16^

However, if the measured melting enthalpy of the ligand-bound protein does change with increased ligand concentrations then (as has been observed)^15^ this is an indication that the equilibrium thermodynamic stability of the native protein structure has also changed due to ligand binding. Therefore, observed changes in thermal stability also reflect directly on thermodynamic consequences of structural (perhaps subtle or not so subtle) perturbations of the protein conformation and stability associated with ligand binding.

Depending on the type and strength of binding interactions, temperature shifts on thermograms of proteins in the presence of ligands can be dramatic (easily tens of degrees). These “T-shifts” along with increases in the melting enthalpies of drug/protein complexes that are associated with binding of ligand can be far more significant than detectable variances in either mass or charge that accompany binding. Consequently, compared to other standard methods, DSC derived thermograms can be more sensitive to binding interactions.

DSC is a well-known physical technique commonly used for detailed quantitative analysis of protein stability. Thus, it is surprising that relatively little attention has been given to application of DSC for analysis of drug/protein interactions. This because previous attempts to utilize DSC for drug binding analysis have met with limited success.^18–20^ Recently we reported on a fast, reliable and accurate procedure using DSC measurements that enables evaluation of thermodynamic parameters for ligand bound protein. There are a number of favorable features of this analytical approach. These include: (1) ease of sample preparation and experimental execution, (2) small volume of sample required (^~^500 μL), (3) no prior knowledge of binding activity required, (4) short processing time (less than 90 minutes), (5) Fully amenable to automated, high-throughput format for parallel screening applications.

Collective features of the process provide a promising alternative approach with which to interrogate drug/protein interactions, and argue that DSC could prove to be a potentially superior screening alternative.

## MATERIALS AND METHODS

### Protein Samples

Human Serum Albumin (HSA) (≥ 99% pure, Lot number: SLBT8667) was purchased from Sigma Aldrich (St. Louis, MO) and received as lyophilized power.

### Buffers and Reagents

Standard buffer for all experiments contained 150 mM NaCl, 10 mM potassium phosphate, 15 mM sodium citrate, pH = 7.4.

### Solutions of HSA

All protein samples were prepared in standard buffer as stock solutions at a concentration of 1.0 mM and stored at 4°C for at least 24 hours before use. For DSC melting experiments protein samples were 27-28 μM (^~^2 mg/mL) determined spectrophotometrically.^5, 21–24^

### Drug Samples

Drug/ligands for this study were procured from commercial suppliers. Naproxen (NAP), bromocresol green (BCG), ibuprofen (IB), captopril (CAP), thimerosal (TMS), fluorescein (FSC), caffeine (CAF) Digitoxin (DTX), bilirubin mixed isomers (BIL), tetracaine (TET), β-Estradiol (BST), and chloroquine (CQ) were purchased from Sigma Aldrich (St. Louis, MO). Decanoic acid (DCA) was purchased from Acros Organics (Geel, Belgium). Metformin (MET) was from MP Biomedical (Irvine, CA); metoprolol (MEP) from Alfa Aesar (Haverhill, MA); buproprion hydrochloride (BPR) from Tokyo Chemical Industry (Tokyo, Japan); Multihance (MH) and Prohance (PRO) from Bracco (Milan, Italy). Gadavist (GAD) and Magnevist (MAG) were purchased from Bayer (Leverkusen, Germany); Dotarem (DOT) from Guerbet (Villepinte, France); Ablavar (AB) from Lantheus Medical Imaging (Billerica, MA); and Δ9-tetrahydrocanabinol (THC) from Cerilliant (Round Rock, TX). The novel compound DM1157^25–27^ was provided by Professor David Peyton (Portland State University). BP-DOTA(side), BP-DOTA(corner), NBAM-DO3A, and BPAM-DO3A were prepared and provided by Professor Mark Woods (Portland State University).

### Sample Preparation

#### Preparation of Drug Samples

Drug solids were dissolved in nanopure water to create stock solutions. Gadolinium contrast agents were provided from the manufacturer in aqueous solution. THC was a 1mg/mL analytical standard prepared in methanol. Aqueous insoluble drugs were prepared as stock solutions in appropriate solvents; digitoxin in chloroform; ibuprofen, β-estradiol, and decanoic acid in ethanol.

#### Protein/Drug Solutions

Solutions of HSA samples for drug binding experiments contained the drug present at different concentrations. Protein concentration was constant in all mixtures at 27-28 μM. Protein/Drug solutions were prepared by adding the desired amount of drug to the protein solution and incubating at 4°C for 24 hours prior to testing.

#### Drug/Ligand Solubilization Procedure

Stock solutions of drug compounds were pipetted into separate 2mL microcentrifuge tubes to yield the correct molar amounts of drug for a 1mL working (drug/protein) solution. For drugs with reported binding constants, a minimum of eight distinct drug/protein samples were prepared spanning the binding concentration regime. For drugs of unknown binding strength, a logarithmic titration of drug concentrations was utilized with further refinements depending on results.

Microcentrifuge tubes containing different amounts of drug samples dissolved in organic solvent were placed in a vacuum concentrator (Savant SpeedVac SVC100). Using standard vacuum concentrator procedures to evaporate off organic solvent containing the sample, resulted in known specific amounts of solid drug in each tube. To each tube, HSA was added at 28 μM with standard buffer to produce a one mL working solution for measurements.

#### Calorimetry Measurements

DSC melting experiments were performed as recently described ^15, 28, 29^ and summarized below (details reviewed in Supplemental Material). Data collection, reduction and analysis procedures were identical to those recently reported ^15, 28, 29^. DSC melting experiments measure the excess heat capacity, *ΔC_P_*, of the sample as a function of increasing temperature. Plots of *ΔC_P_* versus temperature are commonly termed DSC thermograms. These were measured for HSA molecules in mixtures with the various ligands. At least two thermograms were measured for every sample examined.

Thermodynamic parameters determined from thermograms measured as a function of ligand concentration provided a means for evaluation of the dissociation binding constant, *K_D_*.^15^ The actual process was comprised of four steps.

1. Thermograms of drug/HSA complexes were measured over a variety of drug/HSA concentration ratios. These provided measurements of the melting transition temperature, *T_m_* and the transition enthalpy, *ΔH_cal_* from which *ΔG_cal_*(37°*C*) was determined for ligand/HSA mixtures as a function of ligand concentration.
2. Drug concentrations served as the doses, while *ΔG_cal_*(37°*C*) or *T_m_* were the responses measured for HSA in mixtures with each drug at increasing doses. From these data Dose-Response (D-R) curves were generated in the form of *ΔG_cal_*(37°*C*) or *T_m_* versus the log of the ratio of ligand to protein concentrations on a Molar scale.
3. D-R curves from (2) were fit with versions of the modified Hill equation by varying the binding constant and stoichiometry as adjustable parameters.^15, 30–32^
4. Fits of D-R curves from (3) provided quantitative evaluation of the ligand binding constant, *K_D_*.
5. Comparison of D-R curves constructed using *ΔG_cal_*(37°*C*) and *T_m_* from (4) provided semi-quantitative estimates of the binding stoichiometry, n, for each ligand binding to HSA.

## RESULTS AND DISCUSSION

### Drug/Ligand Samples

Chemical compounds chosen for DSC analysis of HSA binding were selected to be representative of a broad variety of drugs; all with known clinical utility. A few compounds with no reported binding activity, or poor aqueous solubility were also examined. In total, HSA binding reactions of 28 different compounds were analyzed.

### Binding of 28 drugs to HSA

Typical binding data for HSA in the presence of drugs were comprised of DSC thermograms measured for a constant amount of HSA in the presence of increasing amounts of drug. The complete sets of thermograms collected for HSA in the presence of increasing concentrations and D-R curves derived from them for the 28 drugs are given in Supplementary Material.

For drugs that exhibited detectable binding to HSA there were clearly measurable variations of the HSA-bound thermograms with increasing drug concentrations. From these data quantitative values of *T_m_* and thermodynamic parameters *ΔH_cal_*, 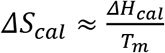, and *ΔG_cal_*(37°*C*) = *ΔH_cal_* − *T* (37°*C*)*ΔS_cal_* were determined at each drug concentration. The values of *ΔH_cal_*, *T_m_* and *ΔG_cal_*(37°*C*) evaluated for thermal denaturation of HSA bound by the 28 drugs are summarized in Table 1 (a more detailed description can be found in Table S1 of Supplementary Material). These results are discussed subsequently.

**Table 1:**
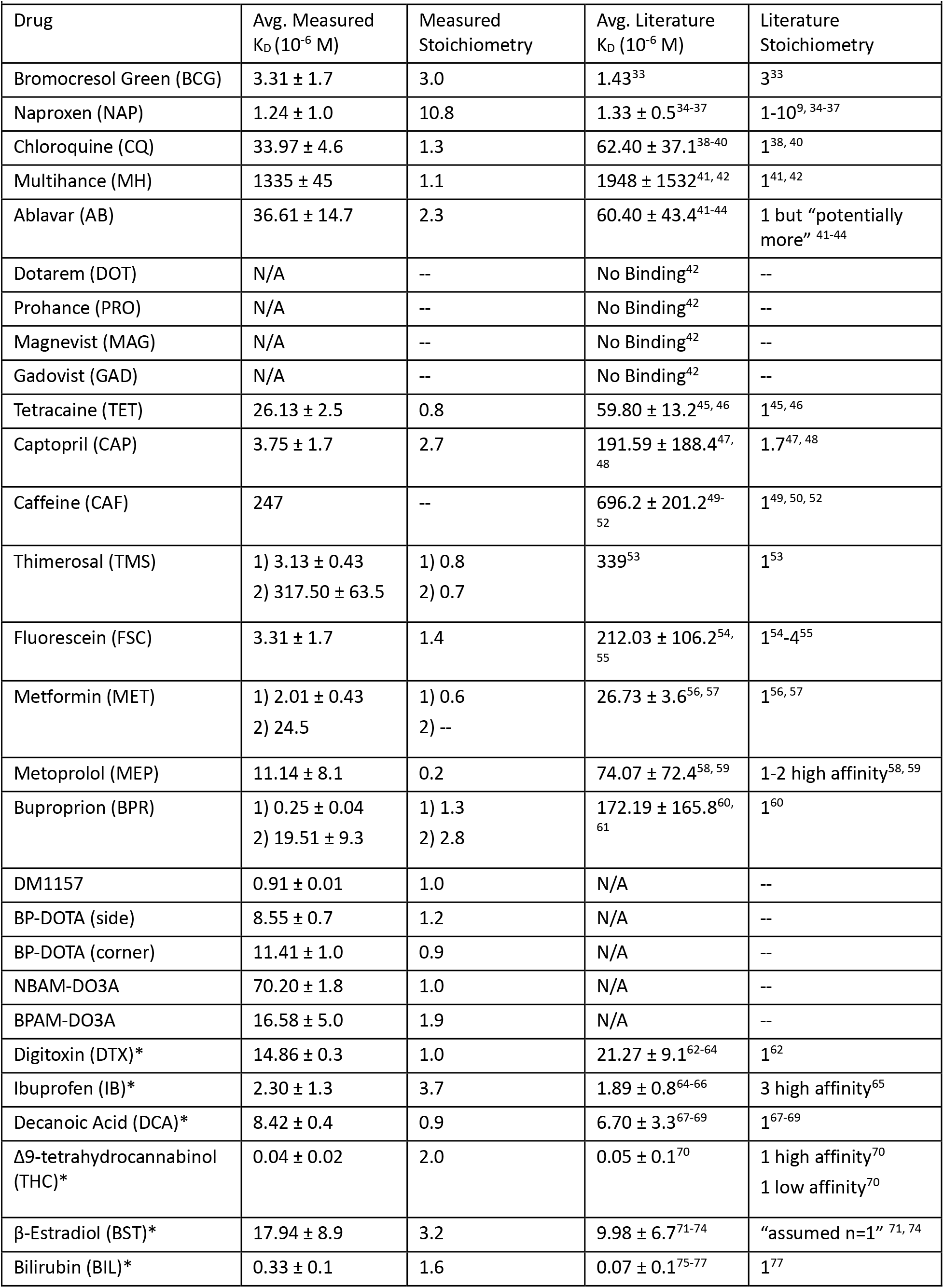
HSA-Drug Binding Constants and Stoichiometries

Out of the 28 drugs examined, 19 of the drugs had binding data reported in the literature. For five of the drugs, no binding data was available. Four of the drugs did not display any binding activity. Evaluated binding constants for the 19 drugs are plotted along with their reported literature values in Fig 1. Evaluated binding constants and stoichiometries for the 28 drugs with HSA are displayed in Table 1 along with their average literature values. Additional information (literature values and experimental techniques for their evaluation) can be found in Supplementary Material. As shown in Fig 1, K_D_ values ranged over five orders of magnitude from 10 nM to 1 mM and agreed with reported values within the error.

**Figure 1:**
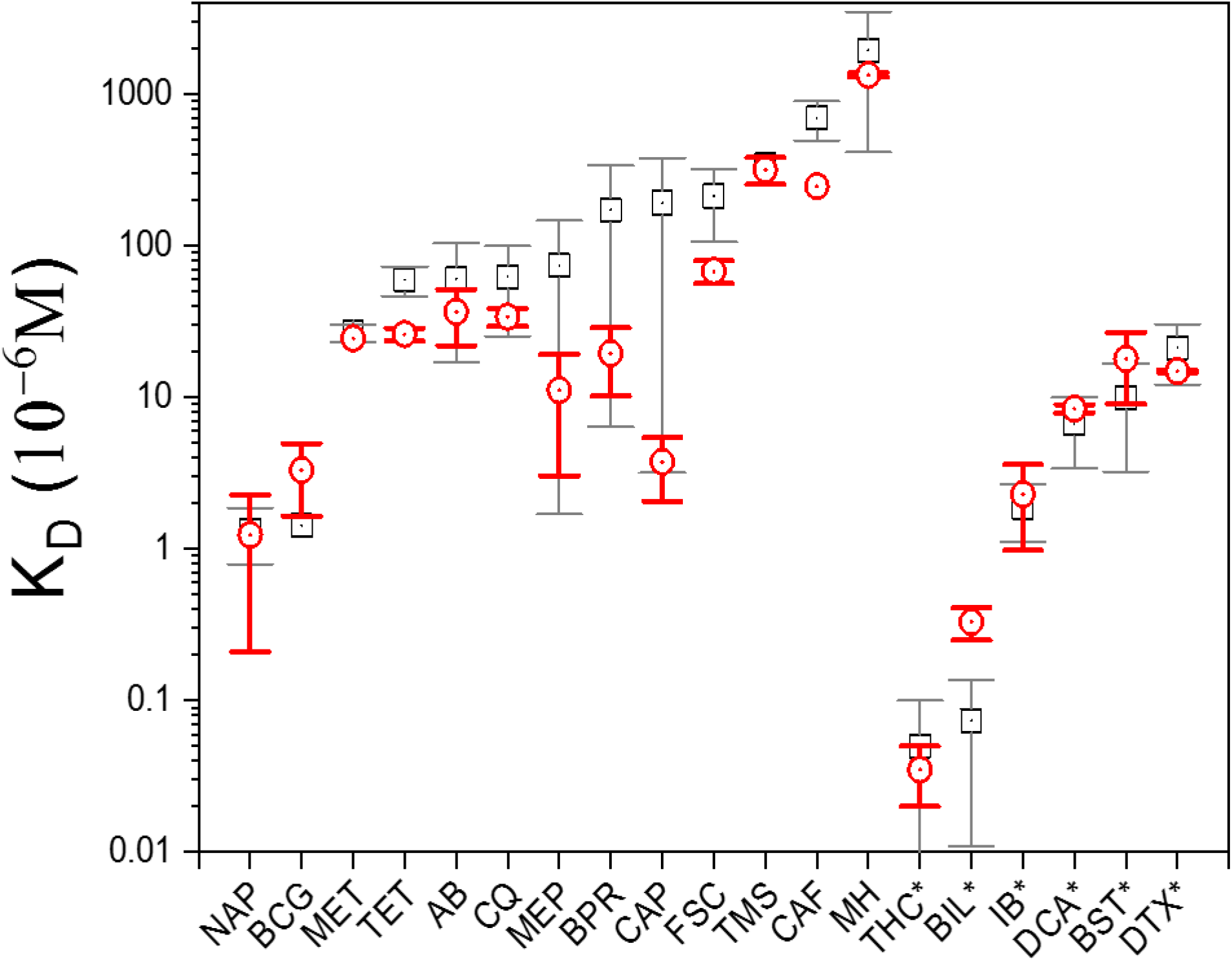
Measured 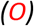 binding constants compared with literature (□). Literature values are given in Table 1. *Insoluble or sparingly aqueous soluble compounds.

To prospect for HSA binding activity for the four drugs with no reported binding activity, thermograms of the drug/HSA mixtures were measured at a drug concentration of 100 mM. This relatively high drug concentration would be expected to evoke some sort of a response in the form of shifted HSA thermograms induced by drug binding. The measured thermograms of these drug mixtures of HSA with 100 mM drug concentration were identical to that of HSA alone, providing no evidence for HSA binding (Supplemental Material).

### Drug/Ligand Binding Constants

For 23 of the drugs, D-R curves constructed with either *ΔG_cal_*(37°*C*) or *T_m_* displayed typical binding behavior where increasing dose concentrations led to a sigmoidal increase in response. These curves were fit well with a single D-R curve and provided evaluations of binding constants in agreement with the literature. Five of the binding curves were atypical and displayed a bi-dose response or destabilization with increased drug concentration. These specific examples are described below.

Two of the drugs (thimerosal and bupropion) displayed bi-D-R curves for both *ΔG_cal_*(37°*C*) and *T_m_*. Analysis of the bi-D-R curves yielded two K_D_ values suggestive of two distinct binding reactions. As shown in Table 1, two binding constants were evaluated for thimerosal, K_D1_ = 3.13 μM and K_D2_= 312 μM (see Table 1). The latter is consistent with the reported literature value, K_D_ = 339 μM ^53^ The former requires additional considerations. One of those is that thimerosal contains an organomercury moiety; with HSA binding likely corresponding to K_D1_. Interestingly, binding of small organomercury compounds to the free sulf-hydryl at cys34 on HSA has been reported to be 0.1-1 μM (with n=4 required for accurate fits). This range of values is very close to our evaluated K_D1_ = 3.13 μM.^78, 79^ Thimerosal rapidly decomposes in the presence of HSA to yield organo-Hg(II) fragments that can also bind HSA.^78^ Detection of two binding reactions for thimerosal is consistent with HSA binding of both whole thimerosal and an un-mercurated thimerosal fragment at its specific site on HSA.^78^

An analogous situation likely exists for bupropion for which the t-butylamino group is reportedly cleaved yielding two products in solution, i.e. a chlorophenyl product and t-butylamino.^80^ The observed bi-D-R curve likely corresponds to separate binding of the two degradation products to HSA, in a manner similar to thimerosal. The result is the observation of two separate binding constants, K_D1_ = 0.25 μM and K_D2_= 19.5 μM (see Table 1). The average of these is 9.8 μM in excellent agreement with the reported HSA binding constant of 6.37 μM.^60^

For metformin, only the *T_m_* D-R curve was bi-phasic. This observation is possibly due to HSA binding of both metformin and a decomposition moiety. Metformin is a biguanide compound. Generally, in aqueous solutions biguanide compounds decompose to urea and other amines.^57, 81^ These amine compounds can bind to HSA, while urea is a known protein denaturant. The bi-D-R curve for *T_m_* displayed a destabilizing trend for HSA as a function of drug concentration, possibly due to urea, while *ΔG_cal_*(37°*C*) displayed a single D-R curve with K_D_ = 2.44 μM. The destabilizing bi-D-R curve for *T_m_* yielded two separate binding constants, K_D1_ = 1.58 μM and K_D2_= 24.52 μM (see Table 1). The latter is in excellent agreement with the reported average value for metformin, K_D_=26.73 μM.^56, 57^ Presumably the former corresponds to binding of amine degradation products. Although consistent with our results this interpretation is by no means conclusive.

Finally, captopril and metoprolol displayed a single D-R curve for *ΔG_cal_*(37°*C*) and a destabilizing D-R curve for *T_m_*. Averages of evaluated binding constants for captopril and metoprolol were K_D_ = 3.75 μM and 11.14 μM, respectively (see Table 1). These binding constants were consistent with the reported literature values.^47, 48, 58, 59^ As discussed above for metformin the presence of urea as a degradation product obviously can destabilize the protein. Origins of the destabilizing D-R curves for captopril and metoprolol are less obvious. For captopril it has been reported that, compared to unbound protein, the observed binding to HSA encourages formation of aggregates at lower temperatures.^82^ This could lead to destabilization of the native HSA structure with increasing captopril concentrations. For metoprolol, although not conclusive, circular dichroism measurements, at increasing concentrations of metoprolol, led to spectroscopic changes interpreted to indicate a decrease in α-helical structure for bound HSA which likely could destabilize the native structure.^58^

For the majority of compounds, quantitative drug binding analysis was performed using simple D-R analysis. For the exceptions described above, quantitatively accurate *K_D_* values were obtained along with additional qualitative insight generated from HSA binding thermograms. These results were encouraging and demonstrated that our DSC approach was able to detect destabilizing behavior due to drug binding, and provide accurate drug binding constants from a single set of experiments. Further, as described below semi-quantitative estimates of the binding stoichiometry were also obtained.

### Binding Stoichiometry

In some cases, differences between the D-R curves constructed using either *ΔG_cal_*(37°*C*) or *T_m_* were observed. As described below semi-quantitative estimates on the ligand binding stoichiometry could be estimated by comparison of the different binding constants evaluated from fitting the *ΔG_cal_*(37°*C*) and *T_m_* D-R curves. For single site binding, stoichiometry n=1, binding constants evaluated using either *ΔG_cal_*(37°*C*) or *T_m_* were in quantitative agreement. However, for ligands with n > 1, i.e. 2-10, ratios of the binding constants determined from *ΔG_cal_*(37°*C*) or *T_m_* doses differed, semi-quantitatively, by a factor of n. The rationale for this observation follows.

*ΔG_cal_*(37°*C*) was obtained from the measured *ΔH_cal_* and *T_m_*. *ΔH_cal_* is comprised of enthalpy changes of the drug and protein target that accompany binding, and effects of binding on the thermodynamic stability of the protein. This includes intermolecular interactions of different types between drug and target. Also included in *ΔG_cal_*(37°*C*) are changes in entropy due to conformational changes in the drug and target upon binding. Therefore, *ΔG_cal_*(37°*C*) was imminently more sensitive than *T_m_* to individual binding events as well as overall effects of binding on HSA structure. For binding sites able to accommodate more than a single ligand, i.e. with binding stoichiometry n >1, effects of binding each ligand in the site are directly reflected in the evaluated *ΔH_cal_* and consequently *ΔG_cal_*(37°*C*).^15^

For ligand binding where n=1, *T_m_* shifts with increased concentrations of ligands that preferentially bind the intact native protein structure, are a natural occurrence due to Le Chatelier’s principle.^16, 17^ This leads to an increase in *T_m_* of the ligand-bound native structure and agrees with our results for ligands with n=1 stoichiometry.

Conversely, observed behavior for ligands with n > 1 is not simply explained by Le Chatelier’s principle. Accordingly, at high ligand concentrations (above stoichiometric amounts) the release of ligand by denaturation of the protein effects the melting equilibrium. However, at low ligand concentrations (below stoichiometric amounts) prior to occupation of all possible binding sites (in an equilibrium sense), re-distribution of the ligand among available binding subsites within the binding pocket can occur. For example Bromocresol Green has been reported to occupy all three subsites (Ia, Ib, Ic) within Sudlow Site I.^33^

Consequently, for n > 1, ligand binding at the lowest sub-stoichiometric ligand concentrations effectively decreases the actual observed site-specific binding constant for subsequent ligand binding events. In this scenario two competing reactions are operative. The first being primary binding of ligand(s) in the binding pocket. The second being redistribution of the ligand(s) within the pocket. Once ligand concentration is high enough to decrease likelihood of redistribution, and the primary binding reaction dominates the process, then titratable increases in *T_m_* are observed.

An alternative possibility is that the ligand can bind at more than one site with relatively high affinity; but binding at some of these binding sites may not be sufficient to affect the melting transition of ligand-bound HSA. In which case, binding of one (or all) of the available sites, either individually or in concert, may be necessary to affect HSA stability as detected by a shift in *T_m_*. For example, naproxen has up to 10 binding sites on the HSA structure with binding constant values within an order of magnitude of one another.^9^ This was consistent with our observed stoichiometry for naproxen of n = 10.8. Note, that these binding effects need not be exclusive and could occur simultaneously.

Thus, for single ligand binding at a single site, *K_D_* values obtained using either *T_m_* or *ΔG_cal_*(37°*C*) would be expected to be essentially the same (as observed). However, for binding multiple ligands at a single site prior to saturation, measured *T_m_* may be largely unaffected until several binding events have taken place at the site and saturation binding is approached. The result was observation of a shifted D-R curve for *T_m_* (by a factor of n, the binding stoichiometry) compared to those for *ΔG_cal_*(37°*C*). This resulted in differing values for the binding constants obtained from the analysis of *T_m_* or *ΔG_cal_*(37°*C*) D-R curves as a function of ligand concentration. In effect, there were actually two binding constants, i.e. *K*_*D*(*T_m_*)_ and *K*_*D*(*ΔG_cal_*(37°*C*))_ evaluated. The ratio *K*_*D*(*T_m_*)_/*K*_*D*(*ΔG_cal_*(37°*C*))_ provided semi-quantitative estimates of the binding stoichiometry, n. As shown in Table 1 the very good agreement between estimated n values with those reported in the literature for the same ligands supports validity of the above proposition. *K_D_* values derived from *ΔG_cal_*(37°*C*) comprised the ensemble of thermodynamic contributions of ligand binding to HSA. Individual evaluated thermodynamic parameters for binding obtained from thermograms can also yield additional insight and provide a means for further classification of drug interactions with HSA.

### Binding Thermodynamics

Formation of drug/protein complexes requires several specific events. These include: disruption of non-covalent interactions with water for both the drug and target molecule i.e. de-solvation; conformational changes in the target and drug and formation of non-covalent interactions between them required for complexation; exchange, association, or dissociation of ions, protons and small molecules that also accompany drug binding. Each of these events can contribute to binding affinity in different ways, but all manifest in values of binding contributions to the enthalpy Δ*H_B_* and entropy, Δ*S_B_*. In our analysis Δ*H_B_* = Δ*H_HSA·L_* − Δ*H_HSA_*. Δ*H_HSA_* is the melting enthalpy of HSA alone and Δ*H_HSA·L_* = Δ*H_cal_* is the denaturation enthalpy measured for the HSA-ligand complexes. *ΔG_cal_*(37°*C*) for the ligand-bound complexes and HSA alone were derived from values of Δ*H_cal_*, Δ*S_cal_* and *T_m_*. Then *ΔG_B_*(37°*C*) = Δ*G_HSA·L_*(37°*C*) − Δ*G_HSA_*(37°*C*). It should be noted that *ΔG_B_*(37°*C*) does not necessarily equal *ΔG′_B_* = −*R*(310*K*)*lnK_D_*. *ΔG_B_*(37°*C*) contains energetic contributions to binding mentioned above that effect thermal denaturation of the ligand-bound protein. Therefore *ΔG′_B_*(37°*C*) calculated from D-R curves was considered to be the energy only due to ligand binding, not including other thermodynamic effects that might also contribute to the measured *ΔG_B_*(37°*C*). To denote *T_m_* changes caused by ligand binding Δ*T_m_* is used where Δ*T_m_* = *T_m_*(*HSA · L*) − *T_m_*(*HSA*).

Thus, a direct link exists between structural stability of HSA and melting stability of ligand-bound HSA complexes. Our approach involves exploiting this linkage by evaluating binding constants through measurements of the effects of ligand binding on the thermodynamics of ligand-bound protein denaturation. An added benefit of our DSC-based analytical process is the enhanced insight that can be gained into relationships between specific physical (stereochemical) and chemical (functional group) features and their effects on protein binding. In particular, as shown below, dissecting the effects on binding activity of different derivatives with specific modifications of a central compound can prove to be most insightful.

Examination of thermodynamic binding parameters listed in Table 2 provides three examples highlighting how semi-quantitative thermodynamic measurements can inform on specific features that might be considered in drug design choices. These are: (1) BP-DOTA (side) and (corner), (2) NBAM-DO3A and BPAM-DO3A, and (3) chloroquine and DM1157. Selected compounds contain examples of stereoisomers and two functional isomers.

**Table 2:** HSA-Drug Thermodynamic Binding Parameters

| Table 2: HSA-Drug Thermodynamic Binding Parameters |  |  |  |
| --- | --- | --- | --- |
| Drug | $\Delta H_B$ (kcal mol<sup>-1</sup>) | $\Delta G_B$ (kcal mol<sup>-1</sup>) | $\Delta T_m$ (°C) |
| **Test Compounds** |  |  |  |
| bromocresol green | 23.21 | 3.82 | 6.29 |
| naproxen | 44.67 | 5.68 | 5.52 |
| chloroquine | 64.36 | 5.25 | 1.43 |
| Multihance | 49.36 | 4.37 | 1.75 |
| Ablavar | 51.71 | 6.85 | 6.53 |
| tetracaine | 41.68 | 4.37 | 3.16 |
| captopril | 57.44 | 4.56 | -0.41 |
| caffeine | 94.13 | 8.00 | 0.47 |
| thimerosal | 73.93 | 14.30 | 16.67 |
| fluorescein | 62.97 | 8.89 | 7.95 |
| metformin | 52.57 | 4.27 | -0.31 |
| metoprolol | 50.60 | 4.06 | -0.40 |
| bupropion | 44.25 | 3.99 | 0.62 |
| **Unknown Compounds** |  |  |  |
| DM1157 | 60.24 | 4.70 | 0.60 |
| BP-DOTA (side) | 25.18 | 4.05 | 6.10 |
| BP-DOTA (corner) | 50.76 | 7.37 | 7.80 |
| NBAM-DO3A | 35.54 | 3.92 | 4.87 |
| BPAM-DO3A | 71.99 | 7.74 | 4.55 |
| **Insoluble or Poorly Soluble** |  |  |  |
| digitoxin | 32.62 | 4.74 | 3.92 |
| ibuprofen | 119.54 | 23.59 | 19.53 |
| decanoic acid | 147.52 | 30.28 | 24.56 |
| $\Delta 9$-tetrahydrocannabinol | 51.40 | 4.45 | 0.28 |
| β-Estradiol | 47.60 | 4.53 | 0.92 |
| bilirubin | 47.27 | 4.15 | 0.30 |
| **Error** | $\pm 7\%$ | $\pm 0.15$ | $\pm 0.15$ |

First, consider several novel analogs of commercially available gadolinium-based contrast agents used for *in vivo* NMR imaging (generous gifts of Professor Mark Woods and Dr. Lauren Rust). DSC measurements of these compounds provided a quantitative metric by which to gauge effects of different specific chemical modifications of the drugs on HSA binding. Specifically, Dotarem (DOTA) and Prohance (DO3A) have been reported to not bind measurably to HSA.^42^ Our analysis concurred and thermograms of drug/HSA mixtures were identical to HSA alone with no binding detected. In contrast, functional and stereochemical modifications of these compounds conferred appreciable HSA binding activity; seen through unique responses of the measured thermograms of their mixtures with HSA, and corresponding evaluated thermodynamic parameters as described below.

Modified forms, BP-DOTA (side) and (corner) are novel stereoisomers of Dotarem^83^ (generous gift of Professor Mark Woods and Dr. Lauren Rust, Portland State University). For these compounds, the DOTA structure was modified with a biphenyl thiourea functional group.^83^ The primary difference between the side and corner isomers of BP-DOTA is the different regiochemistry of the BP functional substituents on the chelate structure.^83^ Binding data in Table 1 for the BP-DOTA compounds indicated little difference in the average binding constants *K_D_* of 8.6 μM and 11.4 μM, for the side and corner isomers, respectively. However, measured thermodynamic binding parameters were significantly different.

Even though BP-DOTA (side) and BP-DOTA (corner) displayed very similar binding constants for HSA, the enthalpic contributions to binding, Δ*H_B_*, were significantly different at 25.18 kcal/mol for BP-DOTA (side) and 50.76 kcal/mol for BP-DOTA (corner). Likewise, BP-DOTA (corner) displayed greater Δ*G_B_* and Δ*T_m_* than BP-DOTA (side). These significant differences suggested a potentially important role of stereochemical structure on HSA binding of drugs within this compound family. These results highlighted how apparently identical compounds with similar binding constants could display significantly different thermodynamic contributions of drug binding to HSA thermostability. It would be much more difficult to obtain comparable results using other methods (NMR, ITC and SPR) commonly applied to assay ligand/protein binding. Our method provided unique information, i.e. both compounds displayed essentially the same binding constants, but very different thermodynamic contributions of binding to HSA thermostability.

Second, consider NBAM-DO3A and BPAM-DO3A, novel functional isomers of Prohance (unpublished, generous gift of Professor Mark Woods and Dr. Lauren Rust, Portland State University). NBAM-DO3A contains a nitrobenzyl group and BPAM-DO3A contains a biphenyl located at the same position on the chelate (DO3A). In contrast to the BP-DOTA compounds above, these DO3A derivatives displayed significantly different average binding constants. With *K_D_* = 70.2 μM and 16.58 μM for NBAM and BPAM, respectively. Notably, for the weaker binder NBAM, Δ*H_B_* = 35.54 kcal/mol, half the Δ*H_B_* of BPAM at 71.99 kcal/mol. Similarly, Δ*G_B_* for NBAM (3.92 kcal/mol) was approximately 50% of that for BPAM (7.74 kcal/mol). Both compounds had similar Δ*T_m_* values. Evaluated stoichiometries were n=2 for BPAM and n=1 for NBAM. After accounting for differences in estimated stoichiometry for BPAM-DO3A and BP-DOTA, the compounds displayed remarkably similar binding constants. This despite differences in the DOTA and DO3A chelates. Thus, the functional biphenyl modification conferred ^~^11 μM binding to the chelates that otherwise did not bind HSA. In this case the functional derivates (NBAM and BPAM) showed differences in both HSA binding and thermodynamic stability.

Third, consider chloroquine, an antimalarial drug, and DM1157 a novel reversed chloroquine compound consisting of a chloroquine backbone with an attached reversal agent (Generous gift from Professor David Peyton and Dr. Katherine Liebman, Portland State University).^25–27^ Unlike the DOTA and DO3A examples above, chloroquine and DM1157 displayed essentially identical thermodynamic contributions of binding to HSA thermodynamic stability, i.e. Δ*H_B_* = ^~^60 kcal/mol, Δ*G_B_* = ^~^5 kcal/mol, and Δ*T_m_*= ^~^1°C for both compounds. It might be expected given the similar thermodynamic behaviors of the compounds they would display similar binding constants. However, this was not the case. For Chloroquine *K_D_* = 33.97 μM while *K_D_* = 0.91 μM for DM1157. The binding constants differ by almost two orders of magnitude! These results indicated that attachment of the reversal agent conferred a much stronger binding constant compared to the unmodified chloroquine. Even though binding of the bulky reversal agent did not significantly affect thermostability of bound HSA. This was consistent with the general observation for these compounds as ligand size did not necessary correlate with thermodynamic stability.

Of 28 drug compounds examined 22 were reported to be soluble in water. Six were reportedly insoluble. Generally, between 40 and 70% of new chemical entities entering the drug development pipeline potentially face bioavailability issues due to low aqueous solubility.^84^ Difficulties and limitations imposed by poor solubility can compromise binding reactions in aqueous environments where most binding experiments were conducted. As an alternative, binding measurements can be performed in organic solvents but could have potentially deleterious effects on the structure and stability of the protein substrate. The organic environment could also adversely affect the stability of ligand/substrate complexes.^85^ To avoid potential problems associated with poor solubility, a novel sample preparation methodology was invoked. Minimum amounts of drug sample (milligrams) are required with complete avoidance of organic solvents in drug/HSA solutions. The method leverages one of the truly remarkable intrinsic properties of HSA. That is the ability of the protein to accommodate extraordinary levels of ligand binding and actually increase ligand solubility in plasma up to seven times higher than the normal solubility limit.^86^ This ubiquitous binding proclivity of HSA forms the basis for the HSA-assisted sample preparation method that was used. It enabled the solubility of normally insoluble compounds in aqueous mixtures with HSA. Effectiveness of the HSA-mediated solubility process was clearly demonstrated for six poorly aqueous soluble compounds denoted by an asterisk in Fig 1 and Table 1. Evaluated binding constants for these compounds with HSA were in excellent agreement with literature reports. Application of this process expands capabilities of DSC for screening potential drug/ligand candidates.

### Conclusion

A number of techniques have been applied to evaluate protein/ligand binding constants and assess drug-HSA binding. These include equilibrium dialysis, ultrafiltration, absorbance and fluorescence spectroscopy, isothermal titration calorimetry (ITC) and Surface Plasmon Resonance (SPR). Of these, ITC and SPR are most comparable (label-free) with our method.

ITC can provide precise evaluation of binding thermodynamics and binding constants at specific temperatures. Despite the sensitivity of ITC, the experiment is not easily scalable for applications in preclinical drug discovery. ITC requires a skilled practitioner to perform experiments, and large sample volumes at high concentrations. In the standard format ITC is incapable of high-throughput applications. Despite being able to deliver perhaps the most quantitatively accurate binding information, because of the inherent drawbacks, ITC has been sparingly employed for drug screening.^11^ Alternatively, the use of SPR for drug binding assays has become a popular screening technique in the drug development industry.

SPR is a label-free detection scheme purported to provide important information about binding of ligands and small molecules to plasma proteins. In fact, SPR has become a method of some choice to examine equilibrium and kinetics of ligand/drug binding. Although potentially quite informative, SPR has short comings and is not without attendant difficulties. SPR does not provide complete information. For example, SPR cannot supply any insight into binding stoichiometry. Another major deficiency of SPR is that proteins or ligands must be immobilized to a surface. For HSA binding, the protein is affixed using a sulfo-NHS reagent.^87, 88^ As recently reported this reagent was used to modify HSA with NHS-biotin.^15^ Modification of HSA molecules acted to decrease measured ligand binding constants by factors from three to 100.^15^ Although SPR is able to quite accurately detect the presence of ligand binding to surface-bound HSA, effects of modifications of HSA (required for surface attachment) on ligand binding can be difficult to gauge. Thus, it can be difficult to evaluate quantitatively accurate binding constants, for natural proteins, from SPR measurements alone. A comparable label-free alternative that uses natural HSA could provide more quantitatively accurate assessment of ligand binding. Ours is such a method.

In this study we employed our recently reported DSC-based process to evaluate HSA binding parameters for 28 different drugs.^15^ Four of the compounds did not display any detectable binding activity; and six were insoluble in aqueous solutions. Results demonstrated quantitatively accurate evaluations of drug-HSA binding constants and semi-quantitative estimates of binding stoichiometries. For all compounds examined, regardless of aqueous solubility, the near universal agreement with published data for known drug/HSA binding constants attests to the general accuracy and robustness of the approach. Results demonstrated a number of many attractive advantages of our method over more laborious and labor intensive, less accurate, slower and less informative ligand screening techniques such as ITC and SPR. Advantages of the method have the potential to considerably improve the pre-clinical drug screening process. More effective preclinical screening can reduce drug development costs, and lead to production of more successful drugs with lower prices; a major goal of modern healthcare management.

## Acknowledgements

We thank Professors David Peyton, Rob Strongin, Mark Woods and Dr. Laura Rust, (Chemistry Department, Portland State University) for their generous gifts of drug compounds and helpful advice.

