## Supplemental Material for "Evaluation of Drug/Ligand Binding Constants for Human Serum Albumin Using Differential Scanning Calorimetry"

**SUPPLEMENTARY MATERIAL**

**Experimental Protocol:** A CSC Model 6100 Nano II-Differential Scanning Calorimeter (TA Instruments, formerly Calorimetric Sciences Corporation) was used for all measurements. Scanning rates were constant at a rate of 1˚C/min. Buffer baselines used were the average of at least five buffer scans versus water measured from 0 to 100˚C.

DSC data were analyzed using recently published procedures [1, 2] as summarized below. These procedures led to the generation of the primary data comprised of DSC thermograms measured for HSA/Ligand mixtures displayed in part (a) of Figs S1-S25 and as described below formed the basis for further analysis of binding.

Five essential steps comprised the data analysis process.

(1) Excess heat (in μW) of the sample was measured as function of temperature. Plots of the raw μW measurements as a function of temperature were constructed.

(2) The average buffer background scan was subtracted from the raw μW versus T curve of the sample constructed in step 1. This resulted in the buffer-corrected curve of μW versus T for the sample.

(3) The curves from step 2 were normalized for protein concentration (2.0 mg/mL), molecular mass HSA (66.5 kDa), partial specific volume of the protein (0.733 $cm^3$ /g) [32] and sample cell volume (0.3268 mL). This produced the normalized excess heat capacity, $\Delta Cp_N(T)$, versus T curve.

(4) The curves from step 3 were baseline corrected using a 3rd order polynomial fit. For points were used for the polynomial fit. These points were defined by the linear regions before and after the transition. The polynomial fit produced a smooth baseline curve.

(5) The baseline curve from step 4 was subtracted from the normalized $\Delta Cp_N(T)$ versus T curve from step 3. The result was the concentration normalized, buffer and baseline corrected $\Delta Cp_N$-baseline(T) ≡ $\Delta C_P$ versus T curve; the *DSC thermogram.*

Thermograms for HSA bound by ligands were characterized by the following experimentally obtained parameters.

(1) $T_m$, the temperature at the maximum peak height.

(2) $\Delta H_{cal}$, the calorimetric enthalpy, evaluated from the integrated area of the measured thermogram.

(3) $\Delta S_{cal}$, which is closely related to the ratio, $\Delta H_{cal} / T_m$, can be semi-quantitatively evaluated and used to calculate the free-energy at $T = 37°C$, $\Delta G_{cal}(37°C) \approx (\Delta H_{cal} - (310.15)\Delta S_{cal}) \equiv \Delta G^0_{37}$.

(4) Values of $T_m$ and $\Delta G_{37}^0$ as function of concentration were used to construct dose response curves, from which quantitative values of ligand binding constants were determined and estimates of binding stoichiometry were made. At least two thermograms were measured at each concentration for most of the samples. Average standard error on $\Delta H_{cal}$ values was approximately 5%. Replicate experiments produced $\Delta G_{cal}(37°C)$ values within ± 0.2 kcal/mol and $T_m$'s within ± 0.15 °C.

The aforementioned analysis relies on two simplifying assumptions.
(1) melting transitions of ligand-bound HSA occurred in a pseudo two-state manner.
(2) Differences between the initial and final $\Delta C_P$ values on thermograms, $\Delta\Delta C_P$, were effectively zeroed out through baseline subtraction. For the ligands alone $\Delta\Delta C_P \approx 0$ compared to the protein. Due to consistency of the results, negative impact of invoked assumptions was presumably small.

**Dose-Response Curves**: As evaluated from thermogram measurements of HSA/Ligand mixtures, $\Delta G_{cal}(37°C)$ and $T_m$ were the responses to the doses, i.e. increased ratios of ligand/protein concentrations used to create the dose response (D-R) curves shown in Fig S1-S25. D-R curves were fit with modified forms of the Hill equation which provided quantitative estimates of $EC_{50}$ for ligand binding to HSA [3-6]. In a few cases, bi-dose response curves were observed and fits yielded two $EC_{50}$ values. Evaluated $K_D$'s for the different ligands that were examined; and comparison of them with those reported in the literature for the same compounds are shown in Table S1.

| Table S1: HSA-Drug Dissociation Binding Parameters | | | | | | |
|---|---|---|---|---|---|---|
| Drug | Measured $K_D$ ($10^{-6}$ M) from $\Delta G^0_{37}$ | Measured $K_D$ ($10^{-6}$ M) From $T_M$ | Measured Stoichiometry | Literature $K_D$ ($10^{-6}$ M) | Literature Stoichiometry | Literature method |
| **Test Compounds** | | | | | | |
| Bromocresol Green | 1.64 ± 0.86 | 4.97 ± 0.67 | 3.0 | 1.43 | 3 | Spectrophotometry[7] |
| Naproxen | 0.21 ± 0.20 | 2.27 ± 0.26 | 10.8 | 0.12 | 1, 4.1 | Fluorescence Quenching [8] |
| | | | | 0.83 | 6 | Equilibrium Dialysis[9] |
| | | | | 1.82 | Assumed 1 | Circular Dichroism[10] |
| | | | | 2.56 ± 0.04 | Assumed 1 | Fluorescence Quenching[11] |
| | | | | - | 10 for $K_D$ within an order of magnitude. | Crystallography[12] |
| Chloroquine | 29.42 ± 6.62 | 38.52 ± 4.34 | 1.3 | 2.1 | N/A | Equilibrium Dialysis[13] |
| | | | | 55.2 | 1 | Fluorescence Quenching[14] |
| | | | | 129.9 | 1 | Capillary Electrophoresis[15] |
| Multihance | 1290 ± 395 | 1380 ± 618 | 1.1 | 179 | “Set to 1” | Mass Spectrometry[16] |
| | | | | 667 | 1 ± 0.58 | NMR Relaxometry[16, 17] |
| | | | | 5000 | 1 | Ultrafiltration[16] |
| Ablavar | 21.93 ± 2.29 | 51.29 ± 11.25 | 2.3 | 12.5 ± 3.24 | 1 | NMR Relaxometry[18] |
| | | | | 21.7 | 1 | Mass Spectrometry[16, 17] |
| | | | | 147 | 1 | NMR Relaxometry[19] |
| Dotarem | None Detected | | - | - | - | NMR Relaxometry[16] |
| Prohance | None Detected | | - | - | - | NMR Relaxometry[16] |
| Magnevist | None Detected | | - | - | - | NMR Relaxometry[16] |
| Gadovist | None Detected | | - | - | - | NMR Relaxometry[16] |
| Tetracaine | 28.62 ± 7.07 | 23.64 ± 9.42 | 0.8 | 46.60 | 1 | Fluorescence (BSA)[20] |
| | | | | 72.99 ± 8.948 | 1 | Fluorescence Quenching (BSA)[21] |
| Captopril | 2.05 ± 0.23 | 5.44 ± 0.31 (destabilizing) | 2.7 | 3.18 | 1.62 | Fluorescence Quenching[22] |
| | | | | 380 ± 28.1 | 1.72 | Displacement Binding[23] |
| Caffeine | 247 ± 33.34 | ----- | ----- | 83.33 | N/A | Fluorescence Quenching[24] |
| | | | | 389 | 1 | Fluorescence Quenching[25] |

| | | | | | | |
|---|---|---|---|---|---|---|
| | | | | 624.6 | 1 | Fluorescence Quenching[26] |
| | | | | 1075 | "Assumed 1" | Fluorescence Quenching[27] |
| Thimerosal | 1) 3.56 ± 0.27<br>2) 381 ± 28 | 1) 2.70 ± 0.30<br>2) 254 ± 52 | 1) 0.8<br>2) 0.7 | 339 ± 2.28 | 1 | Fluorescence Quenching (BSA)[28] |
| Fluorescein | 56.16 ± 23.63 | 79.82 ± 12.82 | 1.4 | 0.104 ± 0.012 | 1 | Fluorescence Quenching[29] |
| | | | | 329 | 4 | Equilibrium Dialysis[30] |
| | | | | 307 ± 17% | "Could not be determined" | Fluorescence Quenching[30] |
| Metformin | 2.44 ± 0.36 | 1) 1.58 ± 1.25<br>2) 24.52 ± 2.37 (destabilizing) | 0.6 | 23.15 ± 0.107 | 1 | Fluorescence Quenching [31] |
| | | | | 30.30 ± 1.06 | 1 | Fluorescence Quenching (BSA)[32] |
| Metoprolol | 19.24 ± 2.08 | 3.04 ± 4.55 (destabilizing) | 0.2 | 218.81 | 1 | Fluorescence Quenching[33] |
| | | | | 0.91 | 2 high affinity | Equilibrium Dialysis (BSA)[34] |
| | | | | 2.5 | 8 low affinity | Equilibrium Dialysis (BSA)[34] |
| Buproprion | 1) 0.21 ± 0.04<br>2) 10.24 ± 2.82 | 1)0.28 ± 0.15<br>2) 28.78 ± 3.91 | 1) 1.3<br>2) 2.8 | 6.37 ± 0.23 | 1 | Fluorescence Quenching[35] |
| | | | | 338 | N/A | HDM-PAMPA Predictive[36] |
| **Unknown Compounds** | | | | | | |
| DM1157 | 0.90 ± 0.51 | 0.92 ± 0.16 | 1.0 | | | |
| BP-DOTA (side) | 7.89 ± 0.86 | 9.21 ± 0.50 | 1.2 | | | |
| BP-DOTA (corner) | 12.37 ± 0.93 | 10.45 ± 1.25 | 0.9 | | | |
| NBAM-DO3A | 72.04 ± 11.35 | 68.38 ± 10.22 | 1.0 | | | |
| BPAM-DO3A | 11.62 ± 3.75 | 21.54 ± 11.68 | 1.9 | | | |
| **Insoluble or Poorly Soluble** | | | | | | |
| Digitoxin | | 14.56 ± 3.10 | 1.0 | 6.6 ± 0.4 | 1 | Equilibrium Dialysis[37] |

| | | | | | | |
|---|---|---|---|---|---|---|
| | 15.15 ± 1.65 | | | 19.2 ± 0.7 | N/A | Affinity Chromatography[38] |
| | | | | 38 | N/A | Surface Plasmon Resonance[39] |
| Ibuprofen | 3.61 ± 2.84 | 0.98 ± 0.08 | 3.7 | 0.37 | 3 high affinity 5 low | Equilibrium Dialysis[40] |
| | | | | 2.4 | 1.4 high 4.7 low | Affinity Chromatography[41] |
| | | | | 2.9 | N/A | Surface Plasmon Resonance[39] |
| Decanoic Acid | 8.96 ± 0.36 | 7.88 ± 0.33 | 0.9 | 10 | 1 | Equilibrium Dialysis[42] |
| | | | | 10 | 1 | Equilibrium Dialysis[43] |
| | | | | 0.1 | 1 | Spectrophotometrically[44] |
| Δ9-tetrahydrocannabinol | 0.048 ± 0.015 | 0.025 ± 0.005 | 2.0 | ≤ 0.1 | 1 high affinity 1 low affinity | Spectrophotometrically[45] |
| β-Estradiol | 9.02 ± 1.34 | 26.86 ± 7.04 | 3.2 | 0.047 | "assumed n=1 for model" | Fluorescence Quenching[46] |
| | | | | 0.055 ± 0.001 | "assumed n=1" | Isothermal Titration Calorimetry[47] |
| | | | | 11.2 | N/A | Solid Phase Micro-extraction[48] |
| | | | | 28.6 | "considerable evidence for multiple binding sites" | Equilibrium Dialysis[49] |
| Bilirubin | 0.25 ± 0.04 | 0.41 ± 0.30 | 1.6 | 0.007 | N/A | Enzymatic Oxidation[50] |
| | | | | 0.2 | N/A | Fluorescence Quenching[51] |
| | | | | 0.015 | 1 | Fluorescence Quenching[52] |

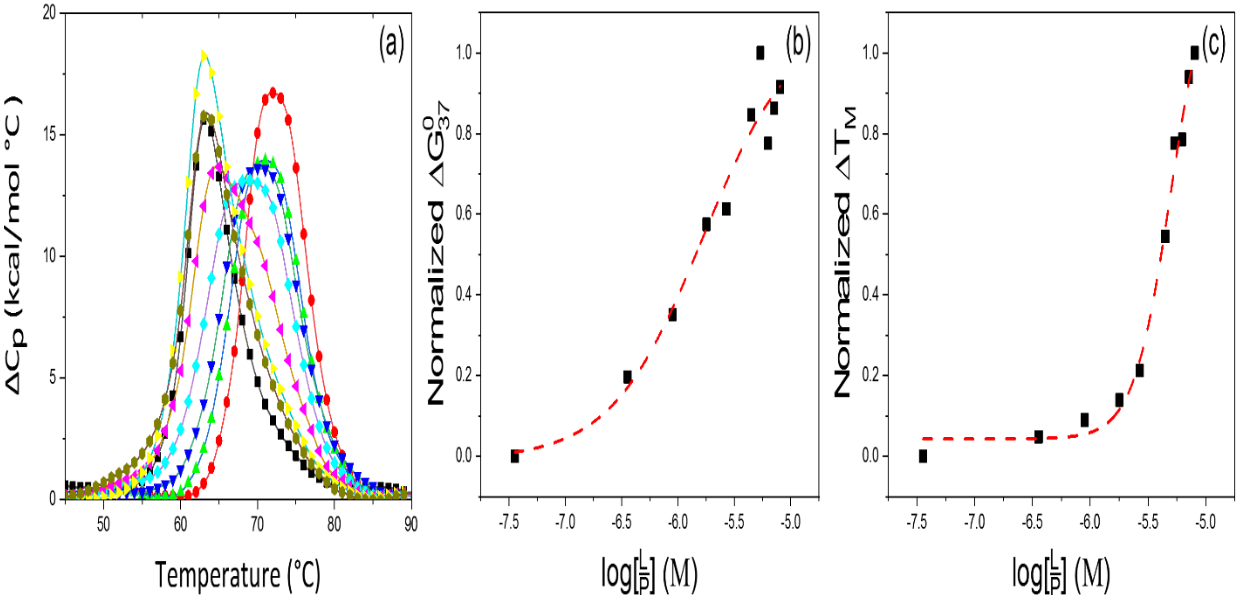


**Figure S1: Binding data for HSA in mixtures with bromocresol green (BCG). (a) Titration thermograms for HSA with BCG as a function of concentration. (■) Standard HSA at 28 μM plus BCG at: 10 μM (⬢), 50 μM (▶), 75 μM (◀), 150 μM (◆), 200 μM (▼), 225 μM (▲), 250 μM (●). (b) Dose response curves constructed from experimentally derived $\Delta G^{0}_{37}$. (c) Dose response curves constructed from experimentally derived $T_M$.**

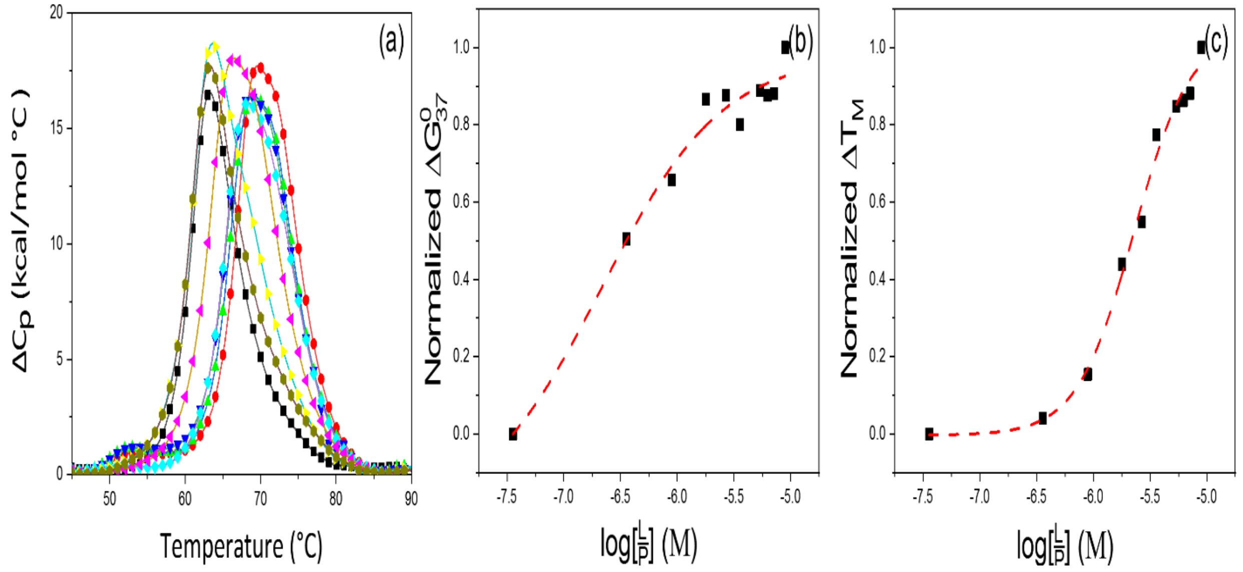


**Figure S2: Binding data for HSA in mixtures with naproxen (NAP). (a) Titration thermograms for HSA with NAP as a function of concentration. (■) Standard HSA at 28 µM plus NAP at: 1 µM (⬢), 10 µM (►), 50 µM (◄), 100 µM (◆), 175 µM (▼), 200 µM (▲), 225 µM (●). (b) Dose response curves constructed from experimentally derived $\Delta G^0_{37}$. (c) Dose response curves constructed from experimentally derived $T_M$.**

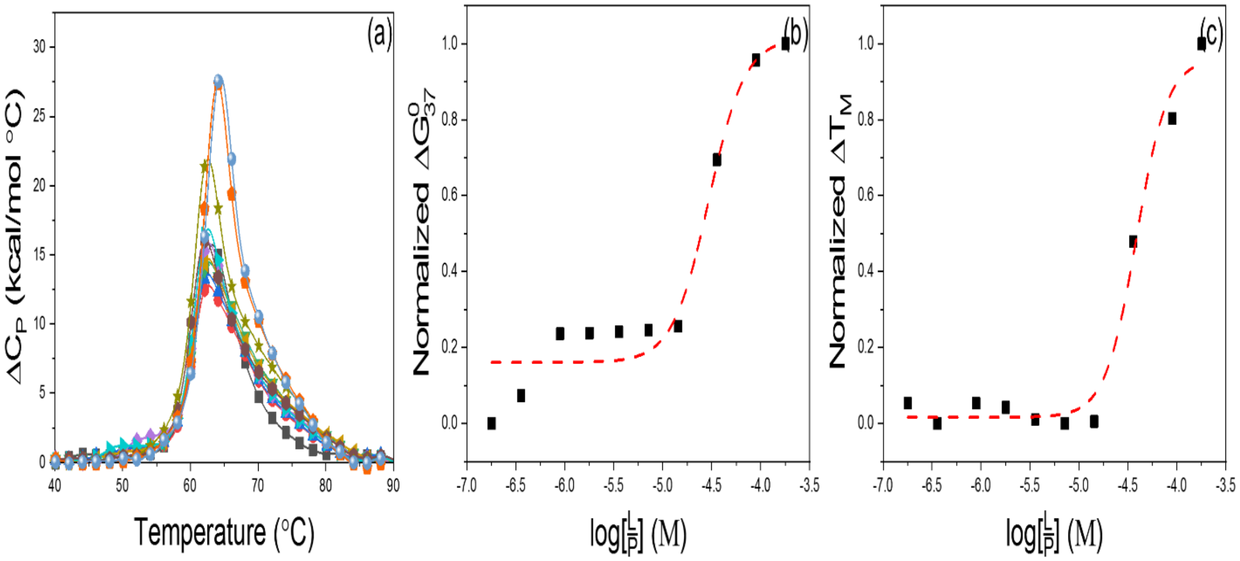


**Figure S3: Binding data for HSA in mixtures with chloroquine (CQ). (a) Titration thermograms for HSA with CQ as a function of concentration. (■) Standard HSA at 2mg/mL plus CQ at: 5 μM (●), 10 μM (▲), 25 μM (▼), 50 μM (◆), 100 μM (◀), 200 μM (▶), 400 μM (⬢), 1,000 μM (★), 2,500 μM (⬟), 4,000 μM (◎). (b) Dose response curves constructed from experimentally derived $\Delta G^0_{37}$. (c) Dose response curves constructed from experimentally derived $T_M$.**

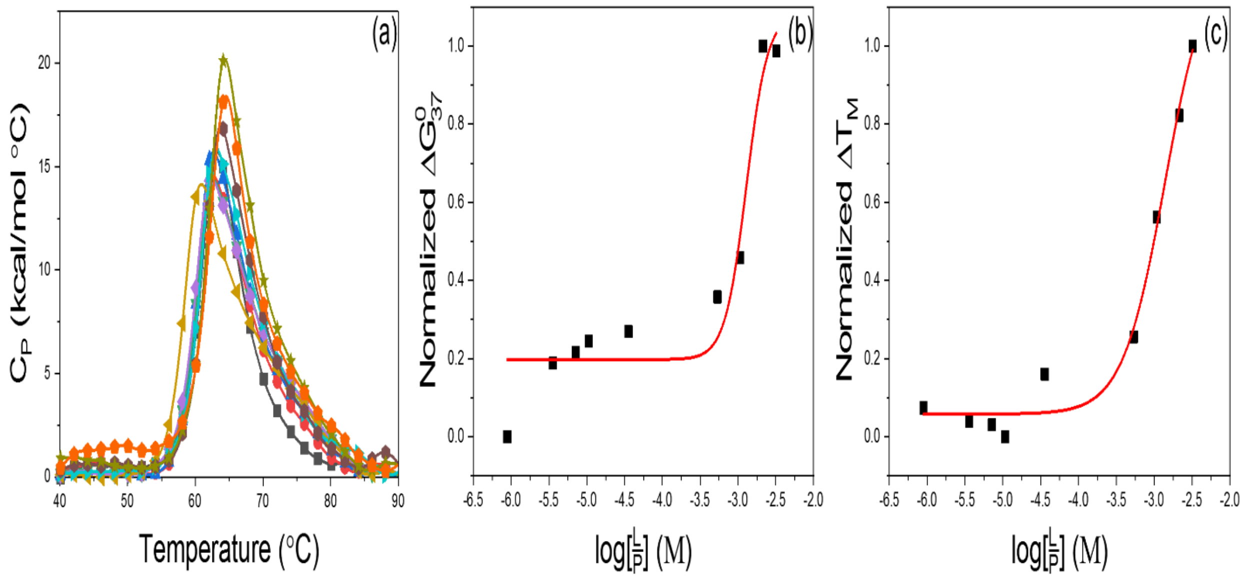


**Figure S4: Binding data for HSA in mixtures with Multihance (MH). (a) Titration thermograms for HSA with MH as a function of concentration. (■) Standard HSA at 2mg/mL plus MH at: 25 μM (●), 100 μM (▲), 200 μM (▼), 300 μM (◆), 1,000 μM (◀), 15,000 μM (▶), 30,000 μM (⬢), 60,000 μM (★), 90,000 μM (⬟). (b) Dose response curves constructed from experimentally derived $\Delta G^0_{37}$. (c) Dose response curves constructed from experimentally derived $T_M$.**

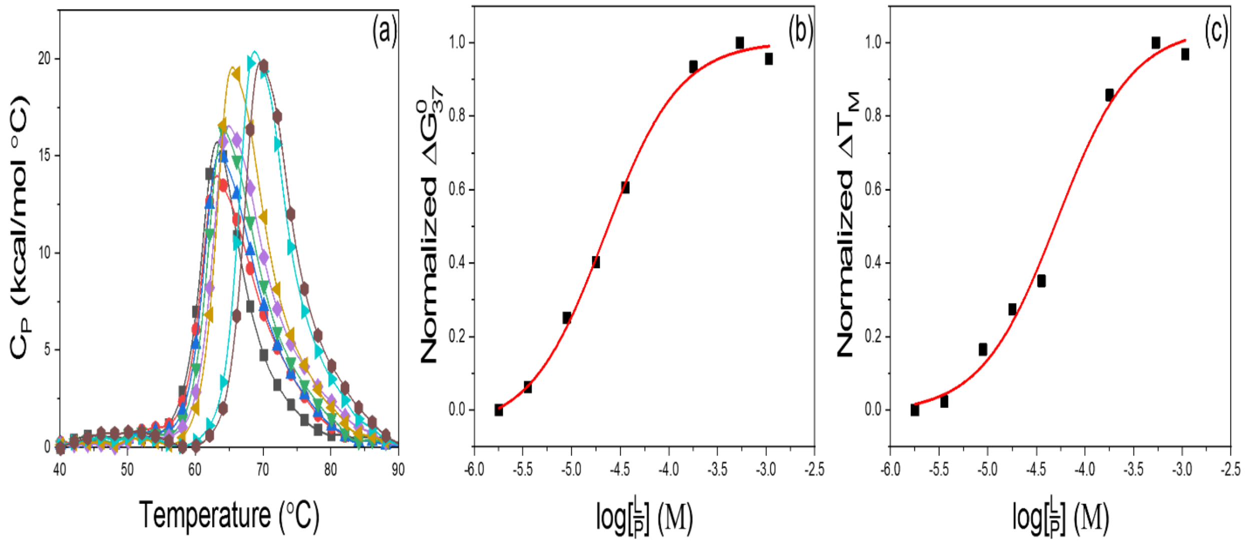


**Figure S5: Binding data for HSA in mixtures with Ablavar (AB). (a) Titration thermograms for HSA with AB as a function of concentration. (■) Standard HSA at 2mg/mL plus AB at: 50 μM (●), 100 μM (▲), 250 μM (▼), 500 μM (◆), 1,000 μM (◀), 5,000 μM (▶), 15,000 μM (⬣). (b) Dose response curves constructed from experimentally derived $\Delta G^0_{37}$. (c) Dose response curves constructed from experimentally derived $T_M$.**

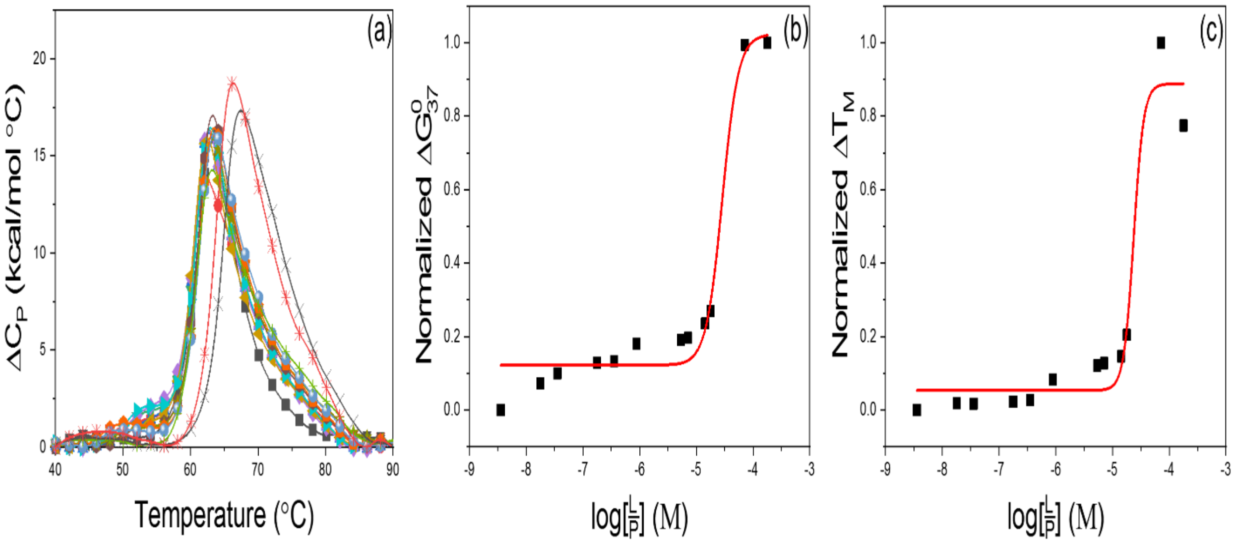


**Figure S6: Binding data for HSA in mixtures with tetracaine (TET). (a) Titration thermograms for HSA with TET as a function of concentration. (■) Standard HSA at 2mg/mL plus TET at: 0.1 μM (●), 0.5 μM (▲), 1 μM (▼), 5 μM (◆), 10 μM (◀), 25 μM (▶), 150 μM (⬣), 200 μM (★), 400 μM (⬟), 500 μM (◎), 1,000 μM (+), 2,000 μM (×), 5,000 μM (✳). (b) Dose response curves constructed from experimentally derived $\Delta G_{37}^{0}$. (c) Dose response curves constructed from experimentally derived $T_M$.**

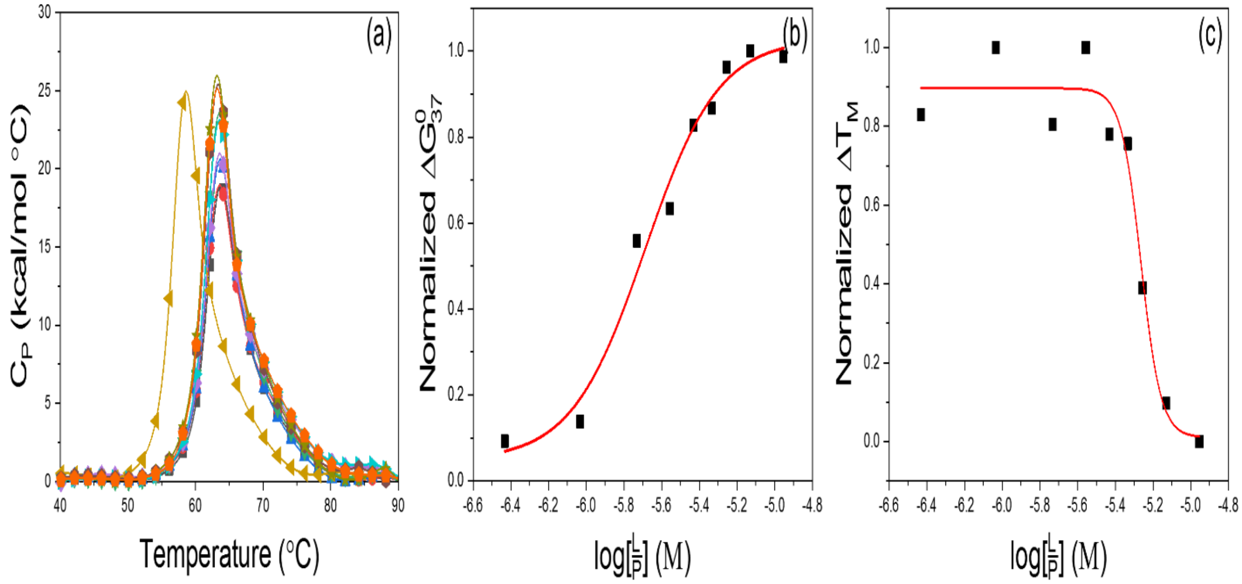


**Figure S7: Binding data for HSA in mixtures with captopril (CAP). (a) Titration thermograms for HSA with CAP as a function of concentration. (■) Standard HSA at 2mg/mL plus CAP at: 10 μM (●), 25 μM (▲), 50 μM (▼), 75 μM (◆), 100 μM (◀), 125 μM (▶), 150 μM (⬣), 200 μM (★), 300 μM (⬟). (b) Dose response curves constructed from experimentally derived $\Delta G^0_{37}$. (c) Dose response curves constructed from experimentally derived $T_M$.**

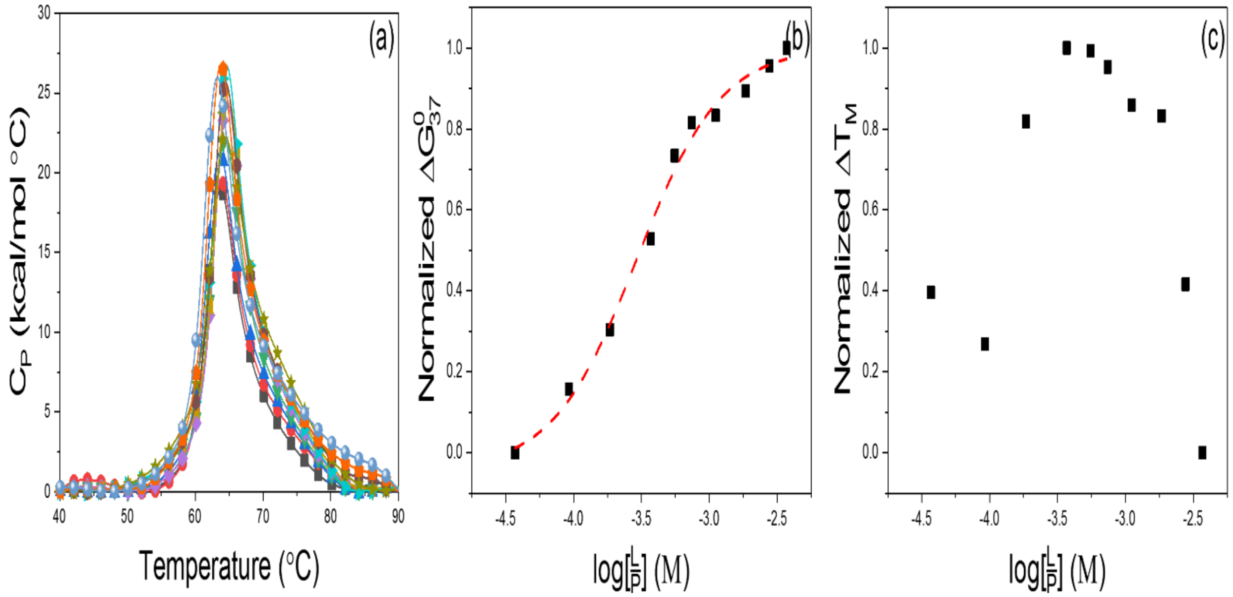


**Figure S8: Binding data for HSA in mixtures with caffeine (CAF). (a) Titration thermograms for HSA with CAF as a function of concentration. (■) Standard HSA at 2mg/mL plus CAF at: 1,000 μM (●), 2,500 μM (▲), 5,000 μM (▼), 10,000 μM (◆), 15,000 μM (◀), 20,000 μM (▶), 30,000 μM (⬢), 50,000 μM (★), 75,000 μM (⬟), 100,000 μM (○). (b) Dose response curves constructed from experimentally derived $\Delta G^0_{37}$. (c) Dose response curves constructed from experimentally derived $T_M$.**

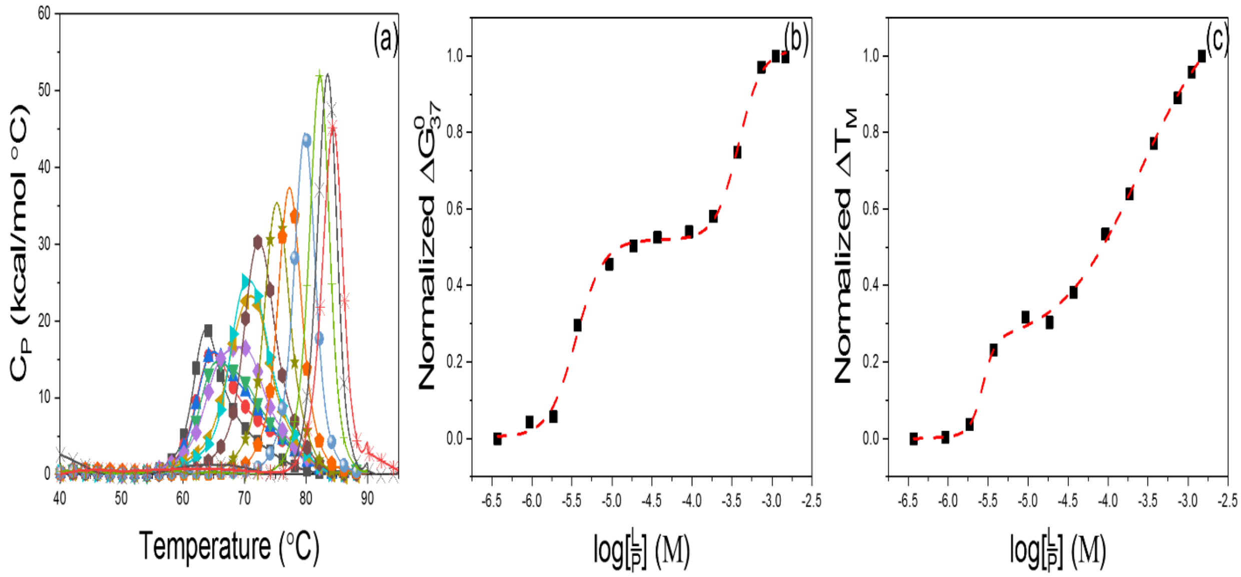


**Figure S9: Binding data for HSA in mixtures with thimerosal (TMS). (a) Titration thermograms for HSA with TMS as a function of concentration. (■) Standard HSA at 2mg/mL plus TMS at: 10 µM (●), 25 µM (▲), 50 µM (▼), 100 µM (◆), 250 µM (◀), 500 µM (▶), 1,000 µM (⬣), 2,500 µM (★), 5,000 µM (⬟), 10,000 µM (◎), 20,000 µM (+), 30,000 µM (×), 40,000 µM (✳). (b) Dose response curves constructed from experimentally derived $\Delta G^{0}_{37}$. (c) Dose response curves constructed from experimentally derived $T_M$.**

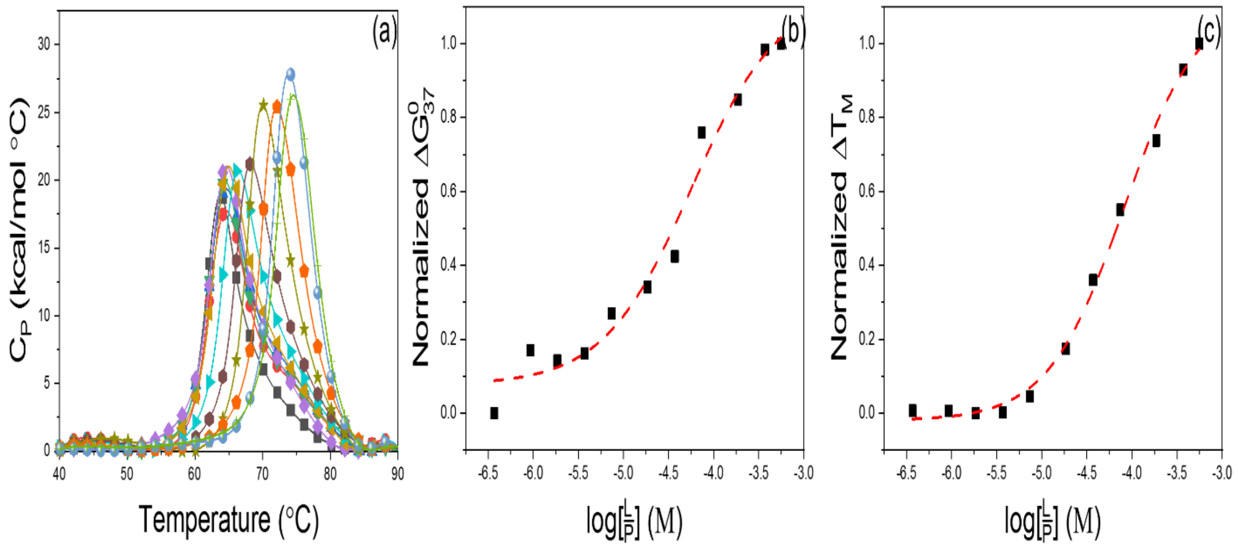


**Figure S10: Binding data for HSA in mixtures with fluorescein (FSC). (a) Titration thermograms for HSA with FSC as a function of concentration. (■) Standard HSA at 2mg/mL plus FSC at: 10 μM (●), 25 μM (▲), 50 μM (▼), 100 μM (◆), 200 μM (◀), 500 μM (▶), 1,000 μM (⬣), 2,000 μM (★), 5,000 μM (⬟), 10,000 μM (◎), 15,000 μM (+). (b) Dose response curves constructed from experimentally derived $\Delta G^0_{37}$. (c) Dose response curves constructed from experimentally derived $T_M$.**

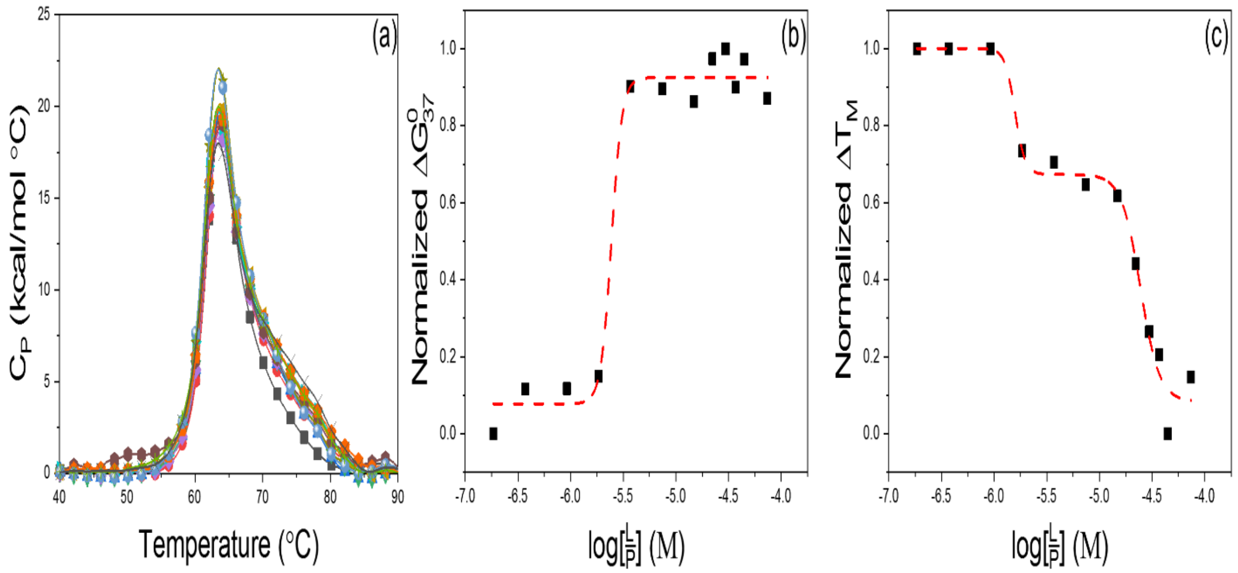


**Figure S11: Binding data for HSA in mixtures with metformin (MET). (a) Titration thermograms for HSA with MET as a function of concentration. (■) Standard HSA at 2mg/mL plus MET at: 5 μM (●), 10 μM (▲), 25 μM (▼), 50 μM (◆), 100 μM (◀), 200 μM (▶), 400 μM (⬣), 600 μM (★), 800 μM (⬟), 1,000 μM (○), 1,200 μM (+), 2,000 μM (×). (b) Dose response curves constructed from experimentally derived $\Delta G^0_{37}$. (c) Dose response curves constructed from experimentally derived $T_M$.**

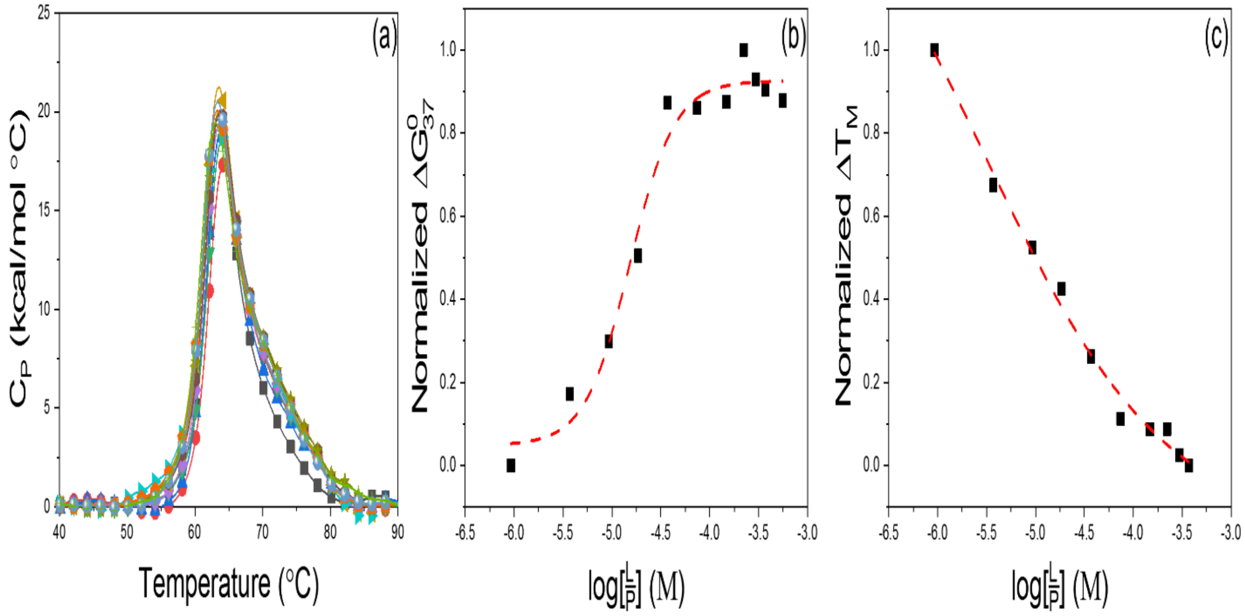


**Figure S12: Binding data for HSA in mixtures with metprolol (MEP). (a) Titration thermograms for HSA with MEP as a function of concentration. (■) Standard HSA at 2mg/mL plus MEP at: 25 μM (●), 100 μM (▲), 250 μM (▼), 500 μM (◆), 1,000 μM (◀), 2,000 μM (▶), 4,000 μM (⬣), 6,000 μM (★), 8,000 μM (⬟), 10,000 μM (◎), 15,000 μM (+). (b) Dose response curves constructed from experimentally derived $\Delta G^0_{37}$. (c) Dose response curves constructed from experimentally derived $T_M$.**

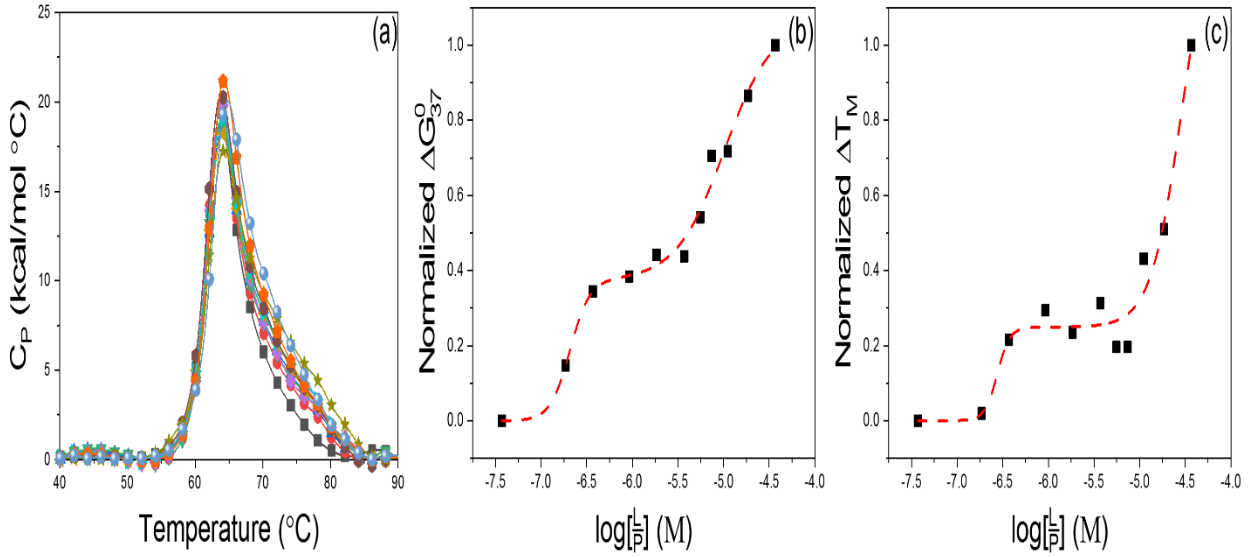


**Figure S13: Binding data for HSA in mixtures with buproprion (BPR). (a) Titration thermograms for HSA with BPR as a function of concentration. (■) Standard HSA at 2mg/mL plus BPR at: 5 µM (●), 10 µM (▲), 25 µM (▼), 50 µM (◆), 100 µM (◀), 150 µM (▶), 200 µM (⬢), 300 µM (★), 500 µM (⬟), 1,000 µM (◎). (b) Dose response curves constructed from experimentally derived $\Delta G^0_{37}$. (c) Dose response curves constructed from experimentally derived $T_M$.**

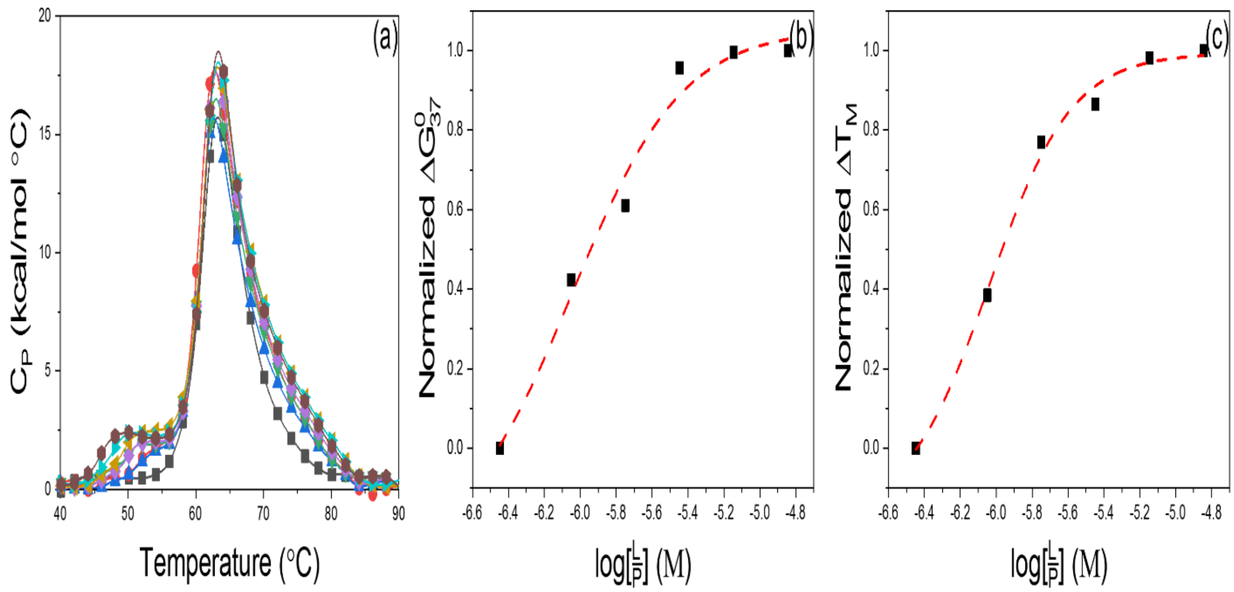


**Figure S14: Binding data for HSA in mixtures with DM1157 (DM1). (a) Titration thermograms for HSA with DM1 as a function of concentration. (■) Standard HSA at 2mg/mL plus DM1 at: 5 µM (●), 10 µM (▲), 25 µM (▼), 50 µM (◆), 100 µM (◀), 200 µM (▶), 400 µM (⬢). (b) Dose response curves constructed from experimentally derived $\Delta G^0_{37}$. (c) Dose response curves constructed from experimentally derived $T_M$.**

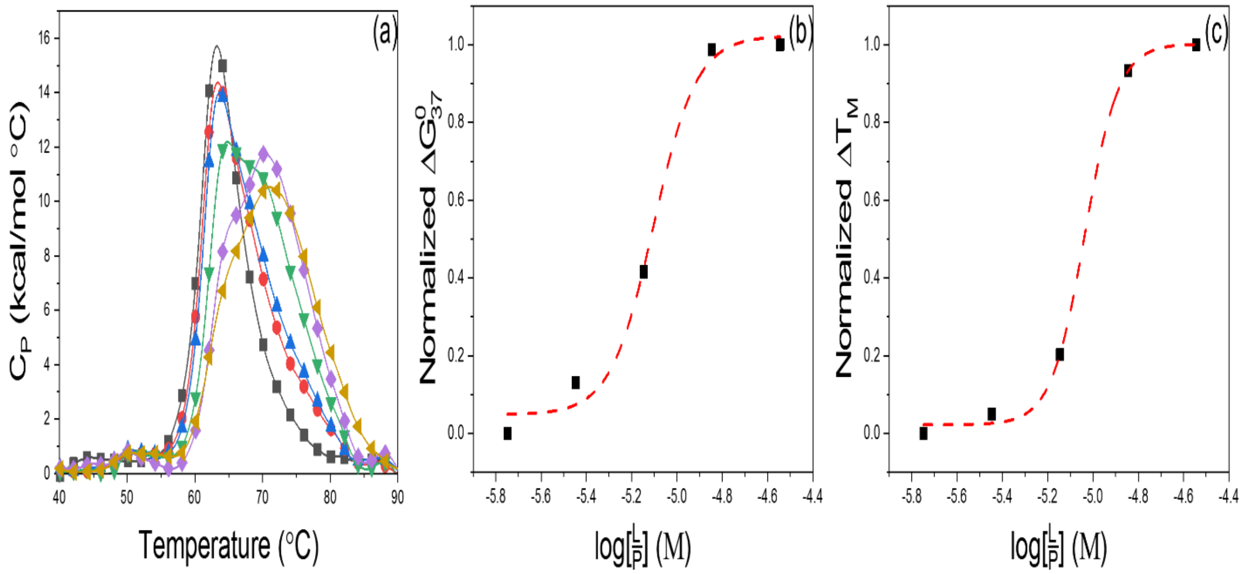


**Figure S15: Binding data for HSA in mixtures with BP-DOTA side (BPs). (a) Titration thermograms for HSA with BPs as a function of concentration. (■) Standard HSA at 2mg/mL plus BPs at: 50 µM (●), 100 µM (▲), 200 µM (▼), 400 µM (◆), 800 µM (◀), 1,000 µM (▶). (b) Dose response curves constructed from experimentally derived $\Delta G^0_{37}$. (c) Dose response curves constructed from experimentally derived $T_M$.**

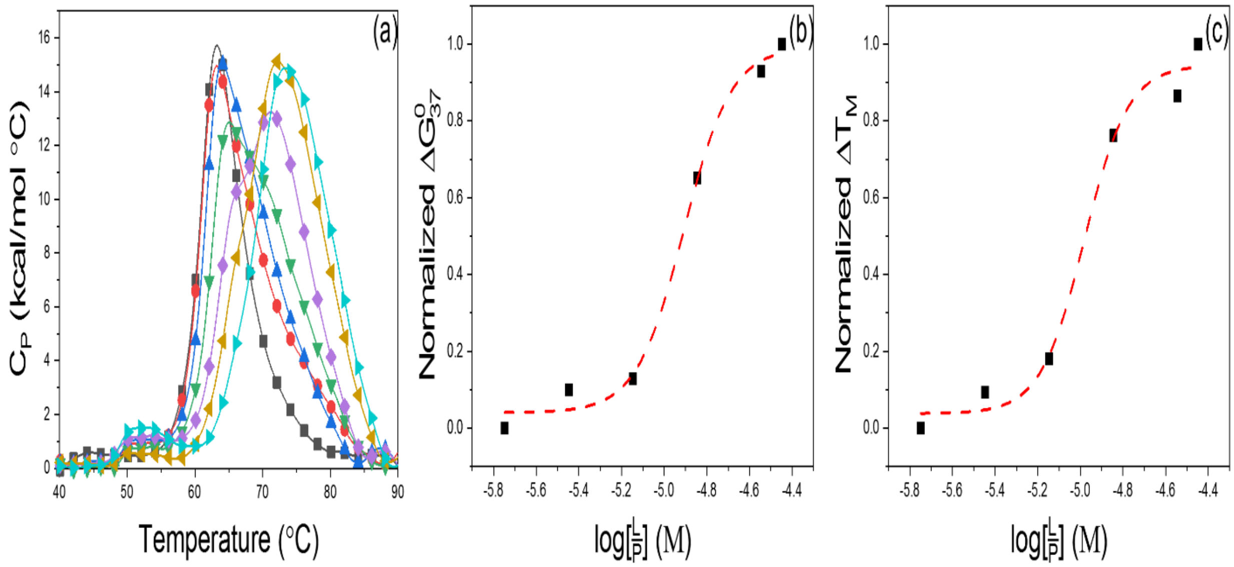


**Figure S16: Binding data for HSA in mixtures with BP-DOTA corner (BPc). (a) Titration thermograms for HSA with BPc as a function of concentration. (■) Standard HSA at 2mg/mL plus BPc at: 50 μM (●), 100 μM (▲), 200 μM (▼), 400 μM (◆), 800 μM (◀), 1,000 μM (▶). (b) Dose response curves constructed from experimentally derived $\Delta G^0_{37}$. (c) Dose response curves constructed from experimentally derived $T_M$.**

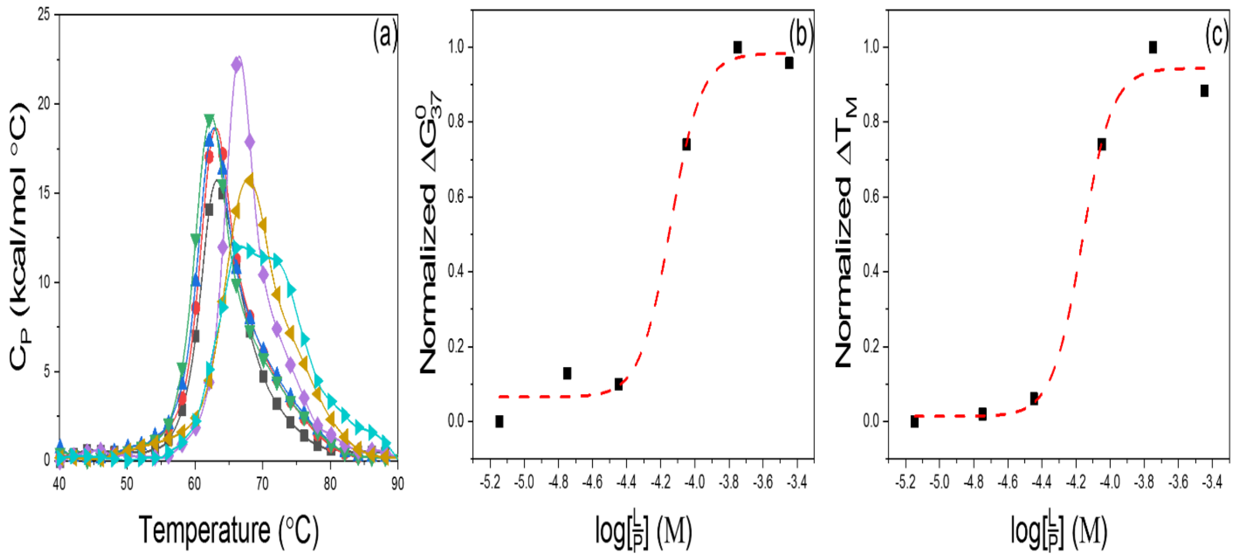


**Figure S17: Binding data for HSA in mixtures with NBAM-DO3A corner (NBD3). (a) Titration thermograms for HSA with NBD3 as a function of concentration. (■) Standard HSA at 2mg/mL plus NBD3 at: 200 μM (●), 500 μM (▲), 1,000 μM (▼), 2,500 μM (◆), 5,000 μM (◀), 10,000 μM (▶). (b) Dose response curves constructed from experimentally derived $\Delta G^0_{37}$. (c) Dose response curves constructed from experimentally derived $T_M$.**

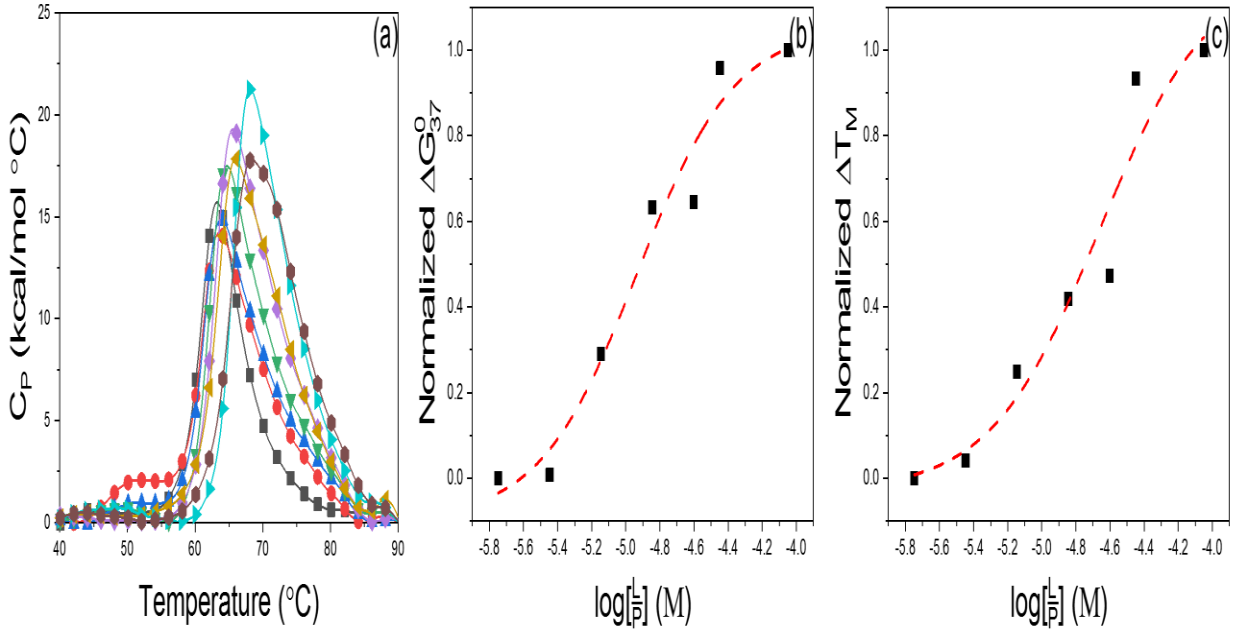


**Figure S18: Binding data for HSA in mixtures with BPAM-DO3A (BPD3). (a) Titration thermograms for HSA with BPD3 as a function of concentration. (■) Standard HSA at 2mg/mL plus BPD3 at: 50 μM (●), 100 μM (▲), 200 μM (▼), 400 μM (◆), 700 μM (◄), 1,000 μM (►), 2,500 μM (⬣). (b) Dose response curves constructed from experimentally derived $\Delta G^0_{37}$. (c) Dose response curves constructed from experimentally derived $T_M$.**

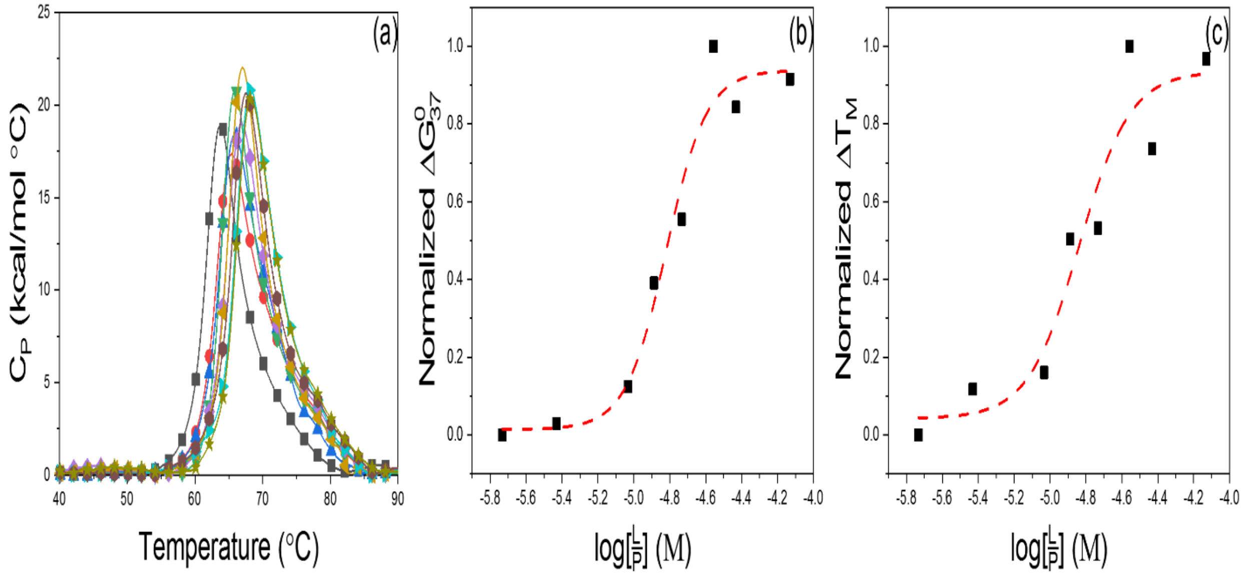


**Figure S19: Binding data for HSA in mixtures with digitoxin (DTX). (a) Titration thermograms for HSA with DTX as a function of concentration. (■) Standard HSA at 2mg/mL plus DTX at: 50 μM (●), 100 μM (▲), 250 μM (▼), 350 μM (◆), 500 μM (◄), 750 μM (►), 1,000 μM (⬣), 2,000 μM (★). (b) Dose response curves constructed from experimentally derived $\Delta G^0_{37}$. (c) Dose response curves constructed from experimentally derived $T_M$.**

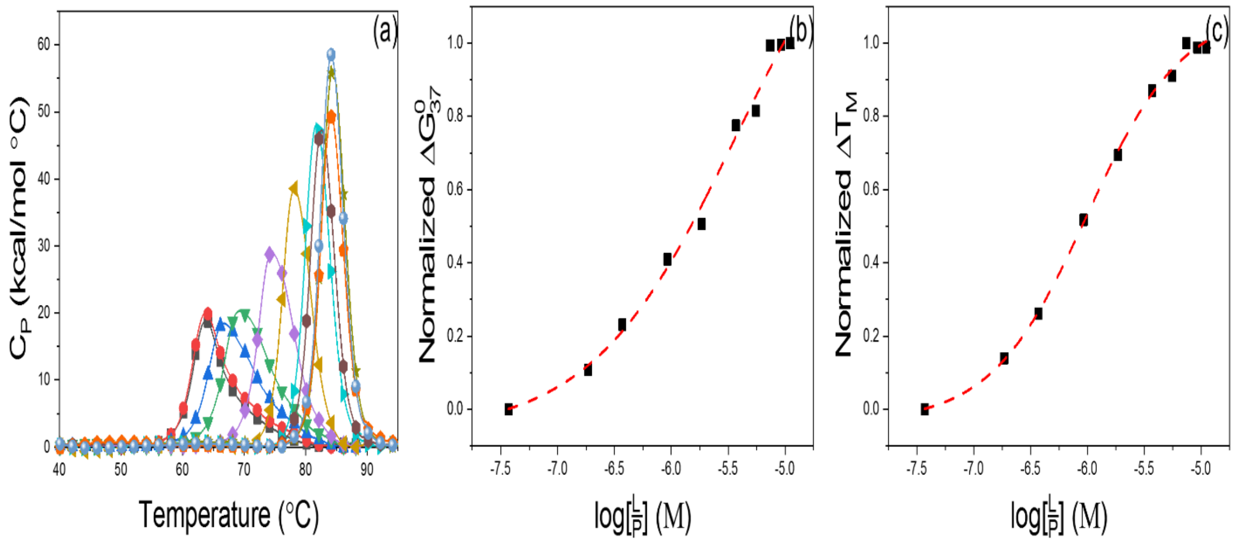


**Figure S20: Binding data for HSA in mixtures with ibuprofen (IB). (a) Titration thermograms for HSA with IB as a function of concentration. (■) Standard HSA at 2mg/mL plus IB at: 1 μM (●), 5 μM (▲), 10 μM (▼), 25 μM (◆), 50 μM (◀), 100 μM (▶), 150 μM (⬣), 200 μM (★), 250 μM (⬟), 300 μM (◎). (b) Dose response curves constructed from experimentally derived $\Delta G^0_{37}$. (c) Dose response curves constructed from experimentally derived $T_M$.**

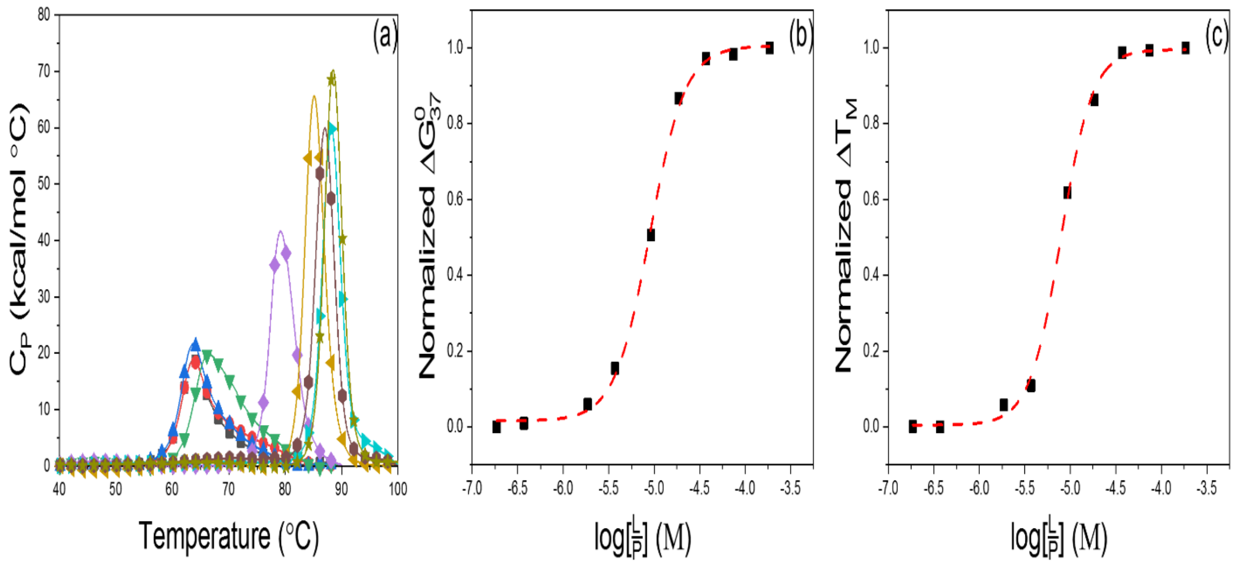


**Figure S21: Binding data for HSA in mixtures with decanoic acid (DEC). (a) Titration thermograms for HSA with DEC as a function of concentration. (■) Standard HSA at 2mg/mL plus DEC at: 5 µM (●), 10 µM (▲), 100 µM (▼), 250 µM (◆), 500 µM (◀), 1,000 µM (▶), 2,000 µM (⬢), 5,000 µM (★). (b) Dose response curves constructed from experimentally derived $\Delta G^0_{37}$. (c) Dose response curves constructed from experimentally derived $T_M$.**

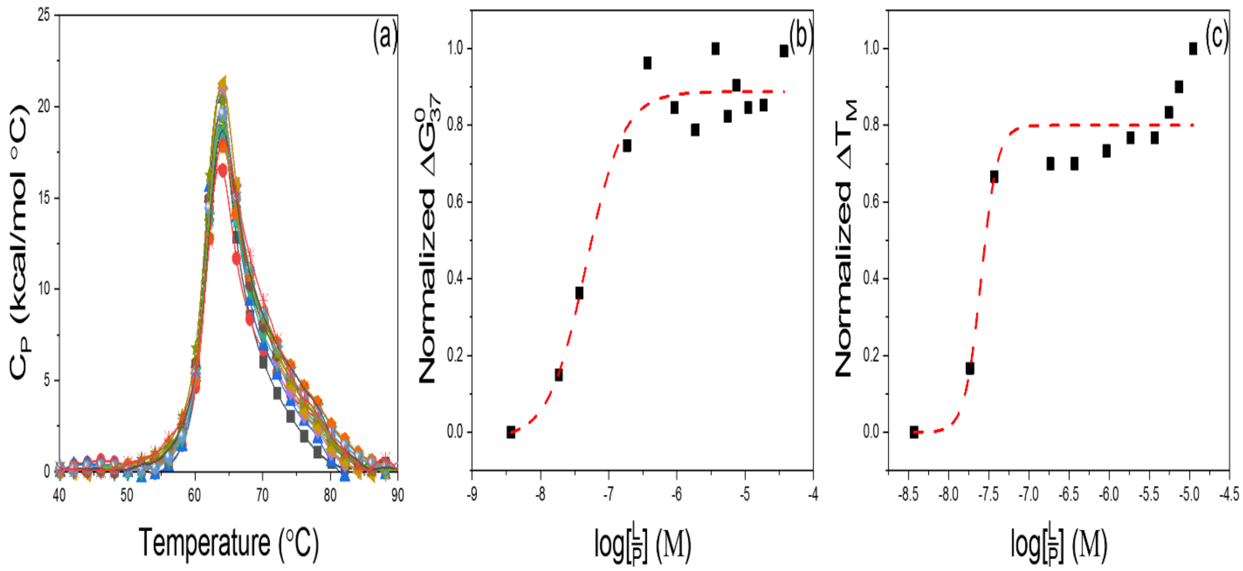


**Figure S22: Binding data for HSA in mixtures with Δ-9-tetrahydrocannabinol (THC). (a) Titration thermograms for HSA with THC as a function of concentration. (■) Standard HSA at 2mg/mL plus THC at: 0.1 μM (●), 0.5 μM (▲), 1 μM (▼), 5 μM (◆), 10 μM (◀), 25 μM (▶), 50 μM (⬢), 100 μM (★), 150 μM (⬟), 200 μM (◎), 300 μM (+), 500 μM (×), 1,000 μM (✳). (b) Dose response curves constructed from experimentally derived $\Delta G^0_{37}$. (c) Dose response curves constructed from experimentally derived $T_M$.**

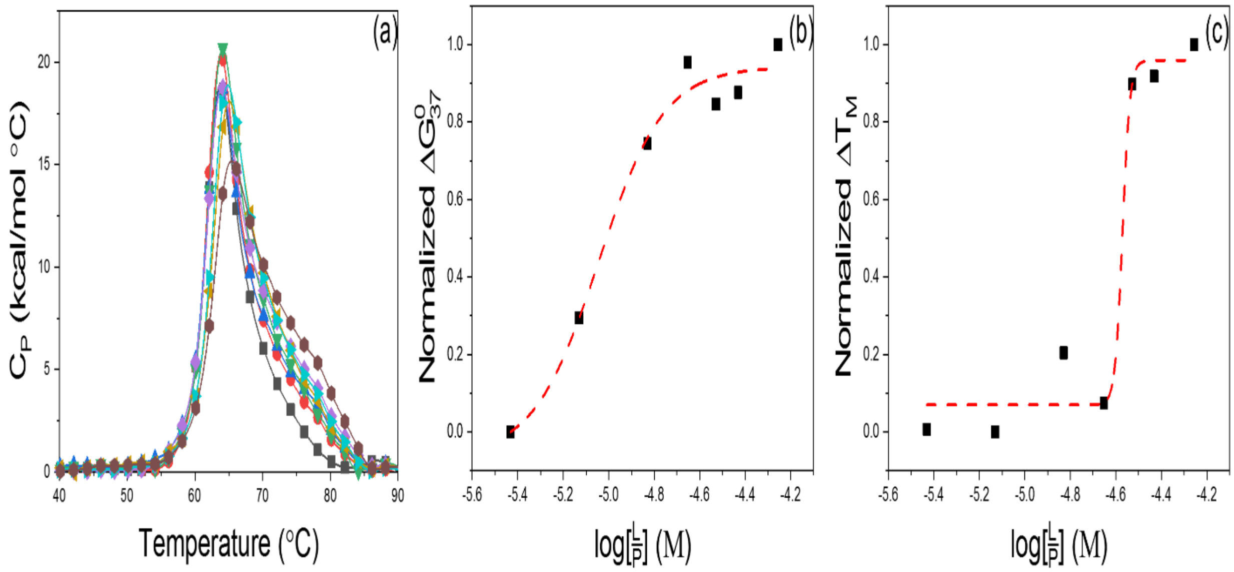


**Figure S23: Binding data for HSA in mixtures with β-estradiol (BST). (a) Titration thermograms for HSA with BST as a function of concentration. (■) Standard HSA at 2mg/mL plus BST at: 100 μM (●), 200 μM (▲), 400 μM (▼), 600 μM (◆), 800 μM (◀), 1,000 μM (▶), 1,500 μM (⬣). (b) Dose response curves constructed from experimentally derived $\Delta G^0_{37}$. (c) Dose response curves constructed from experimentally derived $T_M$.**

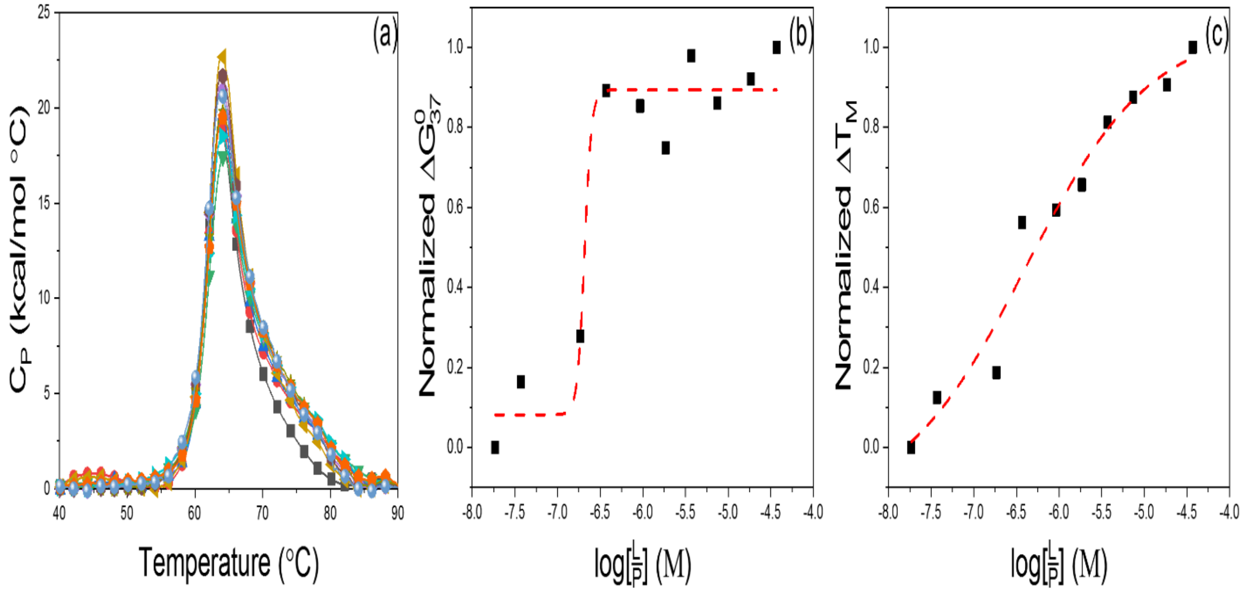


**Figure S24: Binding data for HSA in mixtures with bilirubin (BIL). (a) Titration thermograms for HSA with BIL as a function of concentration. (■) Standard HSA at 2mg/mL plus BIL at: 0.5 µM (●), 1 µM (▲), 5 µM (▼), 10 µM (◆), 25 µM (◀), 50 µM (▶), 100 µM (⬣), 200 µM (★), 500 µM (⬟), 1,000 µM (◎). (b) Dose response curves constructed from experimentally derived $\Delta G^0_{37}$. (c) Dose response curves constructed from experimentally derived $T_M$.**

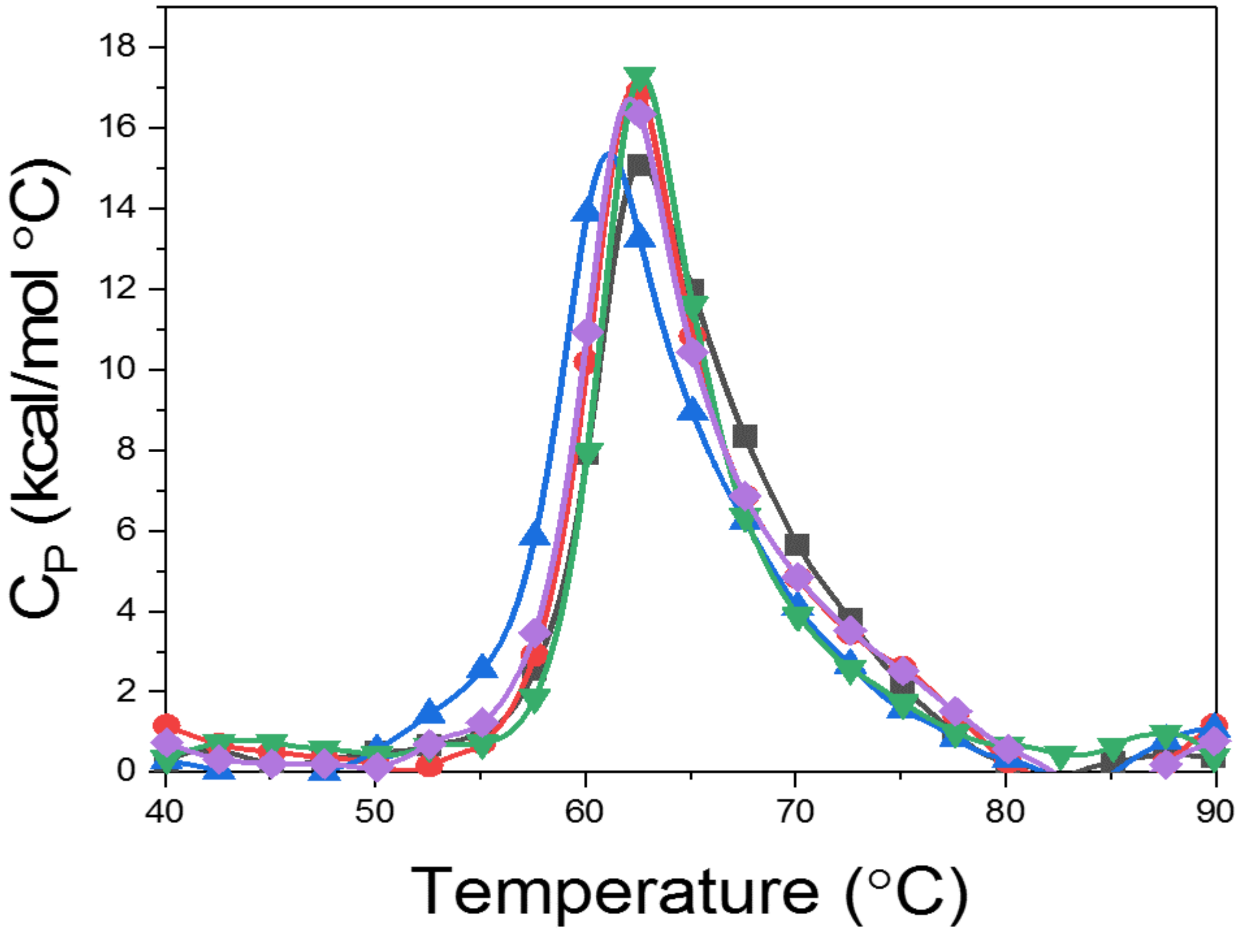


**Figure S25: Thermograms for drugs exhibiting no binding to HSA. Thermograms shown for the compounds are (■) Standard HSA at 2mg/mL with 100,000 µM of (●) Magnevist, (▲) Prohance, (▼) Gadovist, (◆) Dotarem.**
